# Exploring The Role Of Isolation By Distance, Isolation By Environment, And Hybridization In *Campylorhynchus* Wrens Along A Precipitation Gradient In Western Ecuador

**DOI:** 10.1101/2023.08.31.555576

**Authors:** Luis Daniel Montalvo, Rebecca T. Kimball, James Austin, Scott Robinson

**Affiliations:** Florida Museum of Natural History, University of Florida; Department of Biology, University of Florida; Department of Wildlife Ecology and Conservation, University of Florida

## Abstract

Climate variability can cause genetic and phenotypic diversity within species, which affects the evolution of biodiversity. A balance between gene flow and selection maintains changes in the frequency of genetic and phenotypic variants along an environmental gradient. In this study, we investigated a hybrid zone in western Ecuador involving two species of wrens (Aves: Troglodytidae), Campylorhynchus zonatus and C. fasciatus, and their admixed populations. We hypothesized that isolation by distance (IBD) and different ecological preferences, isolation by environment (IBE), result in limited dispersal between populations along the precipitation gradient in western Ecuador.

We asked two main questions: (1) What is the relative contribution of IBD and IBE to patterns of genetic differentiation of these species along the environmental gradient in western Ecuador? And (2) Is there evidence of genetic admixture and introgression between these taxa in western Ecuador? We analyzed 4,409 SNPs from the blood of 112 individuals sequenced using ddRadSeq. The most likely clusters ranged from K=2-4, corresponding to categories defined by geographic origins, known phylogenetics, and physical or ecological constraints. Evidence for IBD was strong across all models, and evidence for IBE was less strong but still significant for annual mean precipitation and precipitation seasonality. We observed gradual changes in genetic admixture between C. f. pallescens and C. zonatus along the environmental gradient. Genetic differentiation of the two populations of C. f. pallescens could be driven by a previously undescribed potential physical barrier near the center of western Ecuador. Lowland habitats in this region may be limited due to the proximity of the Andes to the coastline, limiting dispersal and gene flow, particularly among dry-habitat specialists. We do not propose taxonomic changes, but the admixture observed in C. f. pallescens suggests that this described subspecies could be a hybrid between C. z. brevirostris and C. fasciatus, with different degrees of admixture along western Ecuador and northwestern Peru. This study contributes to the knowledge of avian population genomics in the tropics.

## INTRODUCTION

Geographical barriers are well-established drivers of speciation in many organisms, including birds and mammals (Wright 1943, Tobias et al. 2020, Yao et al. 2022). However, other factors, such as environmental heterogeneity, geographical distance, and hybridization can affect the genetic differentiation of populations (DuBay and Witt 2014, Sexton et al. 2014). The isolation by distance (IBD) and isolation by environment (IBE) models are two common ways to explain how environmental variability and geographic distance affect the genetic structure of populations. IBD occurs when geographic isolation accrues over geographic distance, leading to differentiation (Wright 1943). IBE, on the other hand, is linked to the ability to adapt to environmental factors, which results in strong natural selection when gene flow is low, contributing to the isolation of populations (Alberto et al. 2013, Sexton et al. 2014). The effects of spatial and ecological processes on genetic differentiation are still poorly understood, particularly in tropical species.

The tropics are known to harbor the highest levels of biodiversity, but the true extent of diversity remains underestimated due to incomplete sampling and the lack of detailed studies on many species (Lohman et al. 2010, Bálint et al. 2011). South American bird populations exhibit significant genetic divergence across geographic space, particularly in species complexes, resulting in high intraspecific differences that overlap with interspecies differences (Milá et al. 2009, Tavares et al. 2011, Camargo et al. 2015, Céspedes-Arias et al. 2021, Del-Rio et al. 2022). The Tumbes-Choco-Magdalena biodiversity hotspot in South America, which spans a moisture gradient from the humid Choco-Darien-Western Ecuador region to the dry Tumbesian region, is a region of high-endemism characterized by distinct biogeographical patterns (Dodson and Gentry 1991, Mittermeier et al. 1998). Research suggests that climatic factors might play a role in driving biogeographical patterns, as closely related species’ distributional boundaries match climate regimen boundaries at the transition zone between the two regions in Western Ecuador (Morrone 2006, Albuja et al. 2012, Escribano-Avila et al. 2017, Amador et al. 2019, Prieto-Torres et al. 2019). This suggests that climate may be a major factor influencing the distribution of these species. Western Ecuador provides a unique system for studying genetic structure patterns in which elevation does not vary. However, the factors underlying genetic differentiation in the region remain unknown.

To address this gap, we investigated patterns of genetic structure and the potential contributions of IBD, IBE, and geographical barriers in shaping inter and intraspecific genetic differentiation of two closely related bird species, the Band-backed Wren (*Campylorhynchus zonatus brevirostris*) and Fasciated Wren (*Campylorhynchus fasciatus*). These species exhibit parapatric distributions across wet and dry regions respectively of Western Ecuador and Peru, with the subspecies *C. f. fasciatus* found in the Marañon valley of Northeastern Peru (Figure 1). We hypothesized that different ecological preferences and geographical distances result in limited dispersal between species, and genetic clusters with species along the precipitation gradient in western Ecuador. In the context of testing IBD and IBE shaping distributions of *C. zonatus*, *C. fasciatus*, we asked: What is the relative contribution of IBD and IBE on patterns of genetic differentiation of these species along the environmental gradient in Western Ecuador? If ecological conditions and geographical distance influence genetic differentiation and admixture patterns, then we would expect *C. zonatus* and *C. fasciatus* to show significant associations between genetics, geographical distance (IBD), and/or climate (IBE), leading to population structure that closely mirrors the known geographic distribution of species and populations (Culumber et al. 2012).

**Figure 1.**
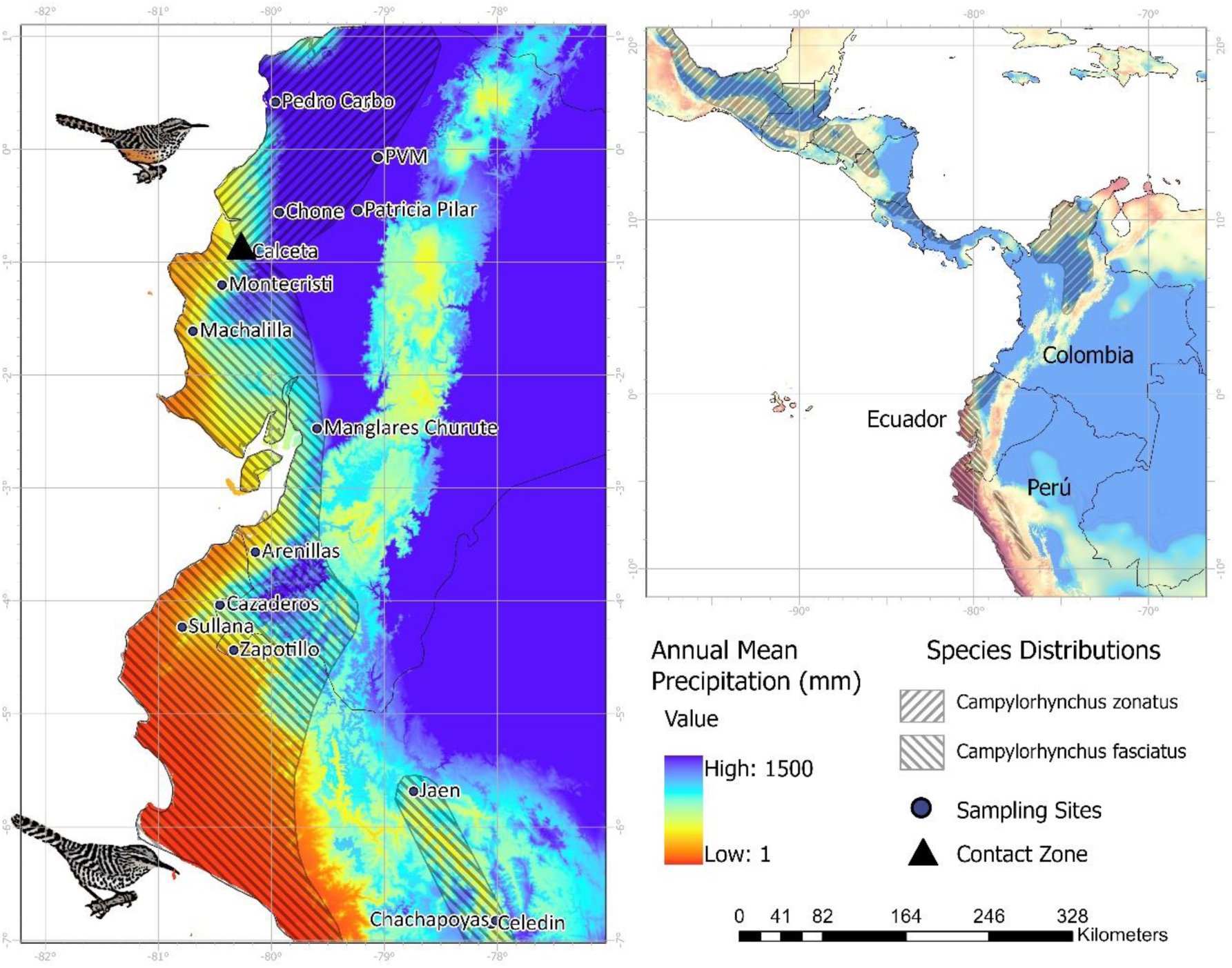
Sampling sites for genomic data and distribution ranges of C. z. brevirostris and C. fasciatus.

Environmental heterogeneity not only promotes genetic differentiation of populations but also facilitates hybridization by providing a pathway for secondary interaction between related species that occur along environmental gradients (Randler 2006, Carling and Thomassen 2012, Runemark et al. 2018). Consequently, hybrid zones, which are regions where genetically distinct populations meet and produce offspring (Harrison and Larson 2014), commonly occur along environmental gradients (Fritsche and Kaltz 2000, Yanchukov et al. 2006, Kameyama et al. 2008). Over 200 avian hybrid zones have been formally described, and environmental gradients have been identified as a key factor in maintaining these hybrid zones (Miller et al. 2014). However, the extent of hybridization can vary depending on the strength of reproductive barriers, ecological factors, and genetic compatibility between the species (Ottenburghs 2018, Winker 2021). Hybrid zones are fundamental for studying evolutionary processes between divergent populations (Whitham et al. 1999, Minder et al. 2007, Sloop et al. 2011).

In addition to the well-established ecological preferences of *C. zonatus brevirostris* and C. fasciatus, field observations of plumage patterns, frequency, and group size suggest that hybridization may occur in the transition zone of the precipitation gradient in western Ecuador (LDM, pers. obs). In the transition zone, some individuals identified as *C. zonatus brevirostris* may lack the ochraceous belly that distinguishes this species (Ridgely and Greenfield 2001, Henry 2005), suggesting that they may be potential hybrids of *C. zonatus* and *C. fasciatus*. Additionally, the frequency and group size of *C. zonatus* increase significantly in this zone (LDM, pers. obs). This suggests that admixed individuals may be present in populations along transitional habitats where *C. zonatus* are locally common. These observations prompted us to ask: Is there evidence of genetic admixture and introgression between these taxa in Western Ecuador? Our study sheds light on the potential role of local adaptation in structuring populations in the region and discusses the implications of physical barriers for the conservation of dry-habitat specialists in Southwest Ecuador and possible hypotheses to explain introgression patterns among species.

## METHODS

### Study Region

The lowland in Western Ecuador and Peru is characterized by a moisture gradient from the humid Choco-Darien-Western Ecuador region in the north to the dry Tumbesian region in the south. Rainfall in Western Ecuador ranges from 2000 to 7000 mm annually, whereas the southern part of the region has an eight-month dry season with less than 1000 mm annually (Dodson and Gentry 1991) (Figure 1).

### Species System

The Band-backed Wren (*Campylorhynchus zonatus*) is found in eastern Mexico to northwest Ecuador, with seven subspecies and four disjunct populations occupying edges, open disturbed areas, and wet forests (Kroodsma and Brewer 2020a). One of its subspecies, *C. z. brevirostris*, is present in two disjunct populations in northwestern Ecuador and northern Colombia. The Fasciated Wren (*Campylorhynchus fasciatus*) is a commonly found species in western Ecuador and northwestern Peru (Figure 1), where it inhabits arid and semiarid areas and deciduous forests (Ridgely and Greenfield 2001, Kroodsma and Brewer 2020b). The subspecies *C. fasciatus pallescens* occurs in western Ecuador and northwestern Peru, while the nominal subspecies *C. f. fasciatus* is present in western Peru and the dry Marañon valley on the eastern side of the Andes (Ridgely and Greenfield 2001, Kroodsma and Brewer 2020b). *C. z. brevirostris* and *C. fasciatus* have parapatric distributions along the precipitation gradient in western Ecuador, with *C. z. brevirostris* restricted to the wet region and *C. fasciatus* to the dry region (Ridgely and Greenfield 2001). Phylogenetic relationships of *Campylorhynchus* show that *C. z. brevirostris* and *C. albobrunneus* are sister species and share a common ancestor with *C. fasciatus* (Barker 2007, Burleigh et al. 2015, Vázquez-Miranda and Barker 2021). Recent studies estimated the time of divergence between *C. z. brevirostris-albobrunneus* and *C. fasciatus-pallescens* at approximately 1.9 million years ago (1.575-2.415 Ma) (Vázquez-Miranda and Barker 2021), whereas the average time to speciation for several phylogroups at the equator is around two million years (Weir and Schluter 2007).

### Samples Collection

We collected blood samples from the brachial vein of 48 putative *C. z. brevirostris* and 49 *C. f. pallescens* individuals at 12 sampling locations along Western Ecuador. Most samples were collected between July and December 2018, with two samples collected in August 2017. We stored the blood samples in Eppendorf tubes with lysis buffer (Tris HCl, pH = 8.0 = 0.1M; NaCl = 0.01M; EDTA = 0.1M; SDS = 3%). We sampled breeding groups at least 400 m apart from one another to prevent capturing related individuals. We obtained tissue samples preserved in 95% ethanol from the Florida Museum of Natural History (FLMNH), including ten samples of *C. f. pallescens* from Sullana in Northwestern Peru (collected in October 2011) and 17 samples of *C. f. fasciatus* from three locations in the Marañon valley of northeastern Peru: Jaen (October 2010), Chachapoyas (November 2009), and Celedin (November 2009). DNA for all samples was extracted using Qiagen DNeasy blood & tissue kits (Qiagen, Germantown, Maryland, USA) and Qiagen extraction protocols. DNA concentration was determined with Qubit dsDNA HS Assay Kit and the Qubit® Fluorometer (Thermo Fisher Scientific Inc., Waltham, Massachusetts, USA).

### Library Construction and Sequencing

We followed the ddRAD-seq protocol by Peterson et al. (2012) and modified by Thrasher et al. (2018). Briefly, we digested 20ul of DNA between concentrations of 3.5ng/ul-40ng/ul with Sbfl and Mspl, and ligated with one of the 20 P1 adapters (each containing a unique inline barcode) and a P2 adapter (P2-Mspl). After digestion and ligation, samples with similar concentrations were pooled in groups of 20 (each with a unique P1 adapter) and purified using 1.5x volumes of homemade MagNA made with Sera-Mag Magnetic Speed-beads (FisherSci) as described by Rohland & Reich (2012). Fragments between 400 bp and 700 bp were selected using BluePippin (Sage Science) by the Cornell University Biotechnology Resource Center (BRC). Following size selection, index groups and Illumina sequencing adapters were added by performing 11 PCR cycles with Phusion DNA Polymerase (NEB). We multiplexed the samples in several index groups (19 and 20 individuals each). These reactions were cleaned up with 0.7x volumes of MagNa and pooled in equimolar ratios to create a single library for sequencing on one Illumina NextSeq 500 (150bp single-end, performed by BRC). The sequencing was performed with a ∼10% PhiX spike-in to introduce diversity to the library.

### Quality Filtering and Demultiplexing

After the quality of the reads was assessed using FASTQC version 0.11.5 (Andrews et al. 2016), we trimmed all sequences to 147bp using fastX_trimmer (FASTX-Toolkit) to exclude low-quality calls near the 3’ of the reads. We removed reads containing at least a single base with a Phred quality score of less than 10 (using fastq_quality_filter). We removed sequences if more than 5% of the bases had a Phred quality score of less than 20. Using the process_radtags module from the STACKS version 2.3 (Catchen et al. 2013), we demultiplexed the reads to obtain files with specific sequences for each individual. After demultiplexing, we retained samples with more than 10^5^ reads for the de novo assembly, removing 12 samples with low read numbers. We ended with a final data set of 112 samples for analysis.

### De novo assembly

Because we do not have a sequenced genome for any species or a close relative, we assembled the sequences de novo using the STACKS pipeline (Catchen et al. 2013). First, we selected 12 samples with the highest number of reads and ran denovo_map.pl testing values from one to nine for -M (number of mismatches allowed between stacks within individuals) and n (number of mismatches permitted between stacks between individuals) following the n=M rule (Paris et al. 2017) while keeping m=3 (stack depth/number of identical reads required to initiate a new allele). We kept r=0.8 (minimum percentage of individuals in a population required to process a locus for that population). It has been shown that at least 80% of the population should present a specific locus to be included, known as the 80% rule or r80 loci (Paris et al. 2017). We set all samples to the same population (p=1) for the parameter testing assembly. We plotted the number of SNPs called against the M parameters to find the optimum M, after which no additional SNP calling was observed. We found an optimum value for n=m=5 for the final de novo assemblies. After the parameters testing assembly, we performed de novo assembly with all the samples and the parameters described above and set p=1. When a RAD locus had more than one SNP, the data were restricted to the first (--write_single_snp) to avoid including SNPs in high linkage disequilibrium (LD). We required a minor allele frequency of at least 0.05 to process a nucleotide site (--min_maf).

We used the dartR package (Gruber et al. 2018) to measure pairwise population-based Linkage Disequilibrium (LD) across all loci. We used 0.5 as the threshold for testing SNPs in LD (Carlson et al. 2004). We retained the entire data set for further analyses, given that only 0.1% of loci showed R2 values over 0.5 across all pairwise combinations.

### Population Structure and Admixture Patterns

We explored visually the genetic structure of the data set with a Spatial Principal Component Analysis (sPCA) analyzed using the function sPCA from the R package Adegenet 2.13 (Jombart 2008). We set the function to build a distance-based connection network with neighbors within a Euclidean distance between one and 26.4km based on the maximum dispersal distance recorded for Cactus Wren (Lynn et al. 2022). The components of the sPCA are separated into global (positive eigenvalues) and local (negative eigenvalues) structures. The global scores indicate either clusters or clines in the dataset compared to between-individual genetic differences reflected by the local scores (Jombart 2008). We assessed the significance of both patterns with a Monte Carlo procedure included in the functions global.rtest and local.rtest using 99,999 permutations.

We used the Bayesian clustering software STRUCTURE version 2.3.4; (Pritchard et al. 2000) to estimate the membership coefficients for each individual (Q-value). We ran a spatial (LOCPRIOR=1) model using sampling locations as prior population information. The model was set to 20 independent replicates with a burn-in of 10^5^ and a run length of 10^6^Monte Carlo iterations —following recommendations by (Gilbert et al. 2012) — for a number of genetic clusters (K) from one to 12, allowing for admixture (NOADMIX=0). We used the method described by (Evanno et al. 2005) implemented in STRUCTURE HARVESTER version 0.6.94; (Earl and Vonholdt 2012) to find the value of K that captures most of the structure in the data, and that seems biologically sensible (Pritchard et al. 2003). We used the software CLUMPP (Jakobsson and Rosenberg 2007) with a LargeKGreedy model and 50000 random repeats to combine replicates accounting for potential “label switching” and “genuine multimodality” differences. We further calculated the posterior probability of assignment of individuals using Discriminant Analysis of Principal Components (DAPC) from the R package Adegenet 2.13 (Jombart 2008). First, we used the function find.clusters to determine the most likely number of genetic clusters and the group membership for each individual using 100 principal components (PCs) and 10^6^ iterations for K=1-20. We selected the number of clusters with the lowest Bayesian Information Criterion (BIC) value as optimal. We used the estimated group membership to perform a preliminary DAPC retaining 100 PCs and two Discriminant Analysis axes (DAs). We used the preliminary DAPC to calculate the optimal number of PCs to keep using the optim.a.score function set with ten simulations. We performed the final DAPCs for K=2-4 using the optimal number of PCs previously estimated.

### Parental and Hybrid Classification

We used the Q-values from STRUCTURE to group individuals as hybrids if 0.1≤Q-value≤0.9 for K=2, and as parental otherwise. We further estimated the Maximum Likelihood of individual Hybrid Indexes (HI, proportion of alleles inherited from one of the parental species). We classified parental individuals if they belong to the northern-most populations surveyed of *C. z. brevirostris* from Ecuador (Las Golondrinas, Pedro Vicente Maldonado, and Pedro Carbo) or the southern-most populations surveyed of “nominally” ssp. *C. f. fasciatus* (Jaen, Chachapoyas, and Celedin) and have Q-values ≥ 0.90 for either parental population, based on STRUCTURE Q-values for K=2. These individuals served as “parentals” to train the function est.h of the R package INTROGRESS 1.2.3 (Gompert and Alex Buerkle 2010). HI ranged from 0 (pure parental *C. z. brevirostris*) to 1 (pure parental *C. f. fasciatus*).

### Genetic Diversity

We analyzed genetic diversity using the genetic clusters defined by STRUCTURE when K=4. DAPC identified K=4 as the most likely number of clusters, while STRUCTURE regarded it as the third most likely. Employing K=4 not only enables a finer classification but also resulted in genetic clusters with geographical boundaries that correspond closely to those of biogeographical regions that have been previously reported (Morrone 2006, Escribano-Avila et al. 2017, Amador et al. 2019, Prieto-Torres et al. 2019) enabling us to explore potential barriers within Western Ecuador.

We assigned each sample to the genetic clusters if their Q-value for that cluster was greater than 0.9. We estimated alleles frequencies, inbreeding coefficient (Fis) per population, and the observed heterozygosity (Ho) per individual using the function gl.report.heterozygosity from the R package dartR (Gruber et al. 2018). We estimated expected heterozygosity (He) as 2*pq*, where *p* and *q* are the proportions of each allele within an individual. We then obtained the 2.5%, 25%, 50%, 75%, and 97.5% percentiles of Ho and He across individuals within genetic clusters. We minimized the potential bias of related individuals in genetic diversity estimates (Jankovic et al. 2010) by selecting samples with low kinship coefficients among birds captured in the same mist net simultaneously. Nei’s Fst estimates and kinship coefficients were estimated with the HierFstat R package (Goudet 2005).

We partitioned the total genotypic variance into components due to differences between genetic clusters and differences between individuals within clusters using analysis of molecular variance (AMOVA) with pairwise Nei’s Fst distances between individuals (Nei and Li 1979). We used the function gl.amova of the dartR package (Gruber et al. 2018) and evaluated levels of significance with 9,999 permutations. Nei’s Fst (Nei and Li 1979) was used to provide a more comprehensive understanding of the genetic differentiation among distinct genetic clusters.

### Isolation by Distance and Environment

We used climate variables from CHELSA 1.2 (Karger et al. 2017). CHELSA is a high-resolution (30 arc sec, ∼ 1km) free global climate data set. We performed a multiple correlation analysis to identify redundancies among the climatic variables using the Hmisc package for R (Harrell 2014). We selected the climatic variables that we considered biologically relevant and had the lowest Pearson correlation coefficients with other selected variables to avoid collinearity. After this process, we selected Annual Mean Temperature (AMT), Annual Mean Precipitation (AMP), and Precipitation Seasonality (PS).

We explored patterns of isolation by distance (IBD) and isolation by environment (IBE) using partial Mantel tests and Generalized Dissimilarity Models (GDM), along with two distinct dissimilarity datasets. The dissimilarity data sets were based on the mean per sampling location of pairwise Nei’s Fst (Nei and Li 1979) and kinship coefficients among samples, normalized as 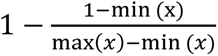 where *x* is the kinship coefficient between two samples. We estimated pairwise Nei’s Fst and kinship coefficients using the HierFstat R package (Goudet 2005). Euclidean distances between coordinates and climate values of each sample were used as predictors of the kinship coefficient matrix. Coordinates and climate values were averaged per sampling location, and then Euclidean distances were estimated and used as predictors for the Nei’s Fst matrix. Geographic and environmental Euclidean distances among the samples were used as predictors for the normalized kinship coefficients. To account for the Andes as a physical barrier, we conducted the partial Mantel tests and GDM using a dataset that included all sampling sites, and another that excluded the eastern sampling sites (Jaen, Chachapoyas, and Celedin).

First, we correlated both dissimilarity matrices against environmental pairwise Euclidian distances controlled by geography using a partial Mantel set up at 9,999 permutations in the R package vegan (Oksanen 2013). We used the log transformation of environmental and geographic distances — suggested for two-dimensional habitats — and *Fst*⁄(1 − *Fst*) for genetic distances following Rousset (1997). Because Mantel test tends to inflate type I error (Guillot and Rousset 2013), we rejected the null hypothesis of no significant correlation if p-value≤0.001 (Diniz-Filho et al. 2013). Next, Generalized Dissimilarity Modeling (GDM; Ferrier and Guisan 2006, Ferrier et al. 2007, Manion 2009) was used to evaluate the association between both genetic dissimilarity datasets as the response variable, and environmental and geographic Euclidian distance as predictor variables. This statistical method uses matrix regression to investigate the relationships between dissimilarities in predictor and response variables, and it has been increasingly used in landscape genetic studies (Freedman et al. 2010, Thomassen et al. 2011, Geue et al. 2016). The GDM model combines multiple matrix regressions (I-splines) into a single non-linear function to analyze how dissimilarity between pairs of sampling locations responds to environmental gradients and geographical distance. In particular, the partial regressions of GDM take into account two important factors: (1) the non-stationary rate of change along an environmental gradient, and (2) the curvilinearity that characterizes the relationship between dissimilarity and environmental gradients (Ferrier et al. 2007, Fitzpatrick et al. 2013). We used the default of three I-splines per predictor. The significance of the model and predictors was tested with 9,999 permutations using the function gdm.varImp of the ’gdm’ package in R (Fitzpatrick et al. 2022). Significance is estimated using the bootstrapped p-value when the predictor has been permuted. The function also estimates the predictor importance measured as the percent decrease in deviance explained between the full model and the deviance explained by a model fit with that predictor permuted (Fitzpatrick et al. 2022).

### Outliers Selection

We searched for possible effects of environmental variables controlled by latitude on potential local adaptations using two different approaches. First, we conducted a multivariate spatial analysis using Moran Spectral Outlier Detection (MSOD) (Wagner et al. 2017). This technique compares the squared correlation coefficients (power spectrum) — between alleles frequencies of each SNP and Moran eigenvectors maps (MEM) — with the average power spectrum of all SNPs (neutral reference). SNPs that deviate over a predetermined threshold from the average power spectrum are selected as outliers capturing the spatial signal of selection. Then, it uses Moran Spectral Randomization (MSR) to test the association between the outlier SNPs and environmental predictors, accounting for spatial autocorrelation (Wagner et al. 2017). Deviation of SNPs from the average power spectrum was measured using z-scores and p-values≤0.01. Significant associations with environmental predictors were tested using 9,999 permutations and p-value≤0.01. We obtained MEM and MSR using the R package Adespatial version 0.3-14 (Dray et al. 2017). Second, we searched for SNP-environmental associations using BAYESCENV version 1.1 (de Villemereuil and Gaggiotti 2015). This Bayesian approach identifies outlier loci — those with large positive Fst values outside of a neutral Fst distribution — significantly correlated with environmental predictors. The model considers population and locus-specific Fst’s, described by a logistic model with a population-specific parameter β, that capture demographic effects; a locus-by-population interaction term γ that reflects the effects of selection of the variable of interest; and a locus-specific term α, that capture the effects of mutations and other forms of selection (de Villemereuil and Gaggiotti 2015). We set the prior for the probability of moving away from the neutral model as π=0.1, and the preference for the parameter α over γ as P=0.5. We used the default parameter settings for the MCMCs. Convergence was tested with Geweke’s, and Heidelberg and Welch’s convergence diagnostics included in the R package Coda (Plummer et al. 2006). Significant effects of the environmental variables were determined using q-values ≤ 0.05 associated with the parameter γ. We regressed climate predictors and latitude to obtain the independent effects of the environmental variables from latitude. We used the average per sampling location of the absolute values of residuals as spatial-independent predictors for BAYESCENV. The environmental values in BAYESCENV should reflect the difference between the observed value in a local population and a reference value (de Villemereuil and Gaggiotti 2015). As all variables are scaled, this already fulfills the requirement of a relative value. The residuals capture the deviations of the ecological variables from the expected values explained by latitude. The mean of absolute values of residuals never exceeded two, which makes them compatible with the default priors for the standard deviation of the model, and standardization was not necessary.

### Gene Ontology

We additionally searched for biological processes (BP), molecular function (MF), and cellular components (CC) associated to the 17 candidate SNPs. We obtained fragment sequences around candidate SNPs from the population.loci.fa file from STACKS. We aligned these sequences to the genome assembly bTaeGut1.4.pri (project GCF_003957565.2) of the zebra finch (*Taeniopygia guttata*) (Formenti et al. 2021). We used the Blast+ command line software (Camacho and Madden 2023) to perform the alignment, with the following parameters: evalue=1 × 10^−20^, word size = 5, and max target seqs = 5000.

We used the chromosome regions of the aligned sequences to obtain gene names according to the HUGO Gene Nomenclature Committee (HGNC) and GO identification codes and using the Ensembl database (Cunningham et al. 2022) and the BioMart software (getBM function; Durinck et al. 2009). We then used the GO identification codes to obtain the gene ontology category and description of the GO term using the bioconductor annotation data package (GO.db; Carlson 2019) and the *Gallus gallus* annotated genome database (org.Gg.eg.db; Carlson 2022). Finally, we used the AnnotationDbi package (Pagès et al. 2023) for R to search for gene functions in GO.db associated with the sequences that included the candidate SNPs.

### Demographic Scenarios

We used Momi2 to examine alternative two-population demographic models that differ in terms of the presence and timing of gene flow between *C. z. brevirostris* and *C. f. pallescens*: (i) pure isolation, (ii) isolation-with-migration, and (iii) isolation with secondary contact (bidirectional and unidirectional in either direction). Because Momi2 models gene flow as pulse events, we inserted four equally distant episodes of gene exchange as a function of divergence time for the isolation-with-migration model. We kept the effective population sizes (Ne) constant within each model. For the secondary contact model, we constrained the migration events to occur after the most recent time boundary of the Last Glacial Maximum (LGM; ∼16,000 years ago) (Heine 2000). We assumed a mutation rate of 4.9 × 10^−9^ substitutions per site per generation (Smeds et al. 2016) and a generation time of two years reported for *Campylorhynchus nuchalis* (Bird et al. 2020). We first performed ten optimizations for each model with ancestral Ne set to 1 × 10^5^ and the stochastic_optimization function to set 100 mini-batches and ten iterations to get initial parameter estimates with reduced computational effort. Based on these results, we used the mean of Ne (4.7 × 10^5^) and time since divergence (1.63 × 10^5^) across runs and models as initial values for subsequent runs. Here, we report parameter values from models with 50 optimizations and initial values as mentioned above and the function stochastic_optimization set with 1000 SNPs per mini-batch and ten iterations. We used the parameter estimations of the 50 runs to generate the mean/median and 95% confidence interval for each estimated demographic parameter. For each model, we selected the optimization with the largest maximum likelihood value for model selection. We used the relative Akaike information criterion to select the best-fit model (Sakamoto et al. 1986). Finally, we assessed the effect of ancestral Ne on model selection by running each model twice as described in step two but with ten optimizations and ten different ancestral Ne ranging from 1 × 10^5^ to 1 × 10^6^.

## RESULTS

After filtering, we obtained 4,409 SNPs with an average coverage per individual of

35.4x (min=12.8x, max=77.9x) and an average genotype call rate of 93.5% from 112 individuals and 16 sampling locations.

### Population Structure and Admixture Patterns

According to the sPCA analysis, the first principal component explained 40% of the variance in the dataset, while the second principal component explained 8.7%. We found significant evidence for global structure in the dataset (p-value<0.001), but not for local structure (p-value=0.474). Upon visually inspecting the SPCA results, we observed two major clusters that corresponded to the species-level distribution of *C. fasciatus* and the subspecies *C. z. brevirostris* in the northwest region of Ecuador (Fig. 2).

**Figure 2.**
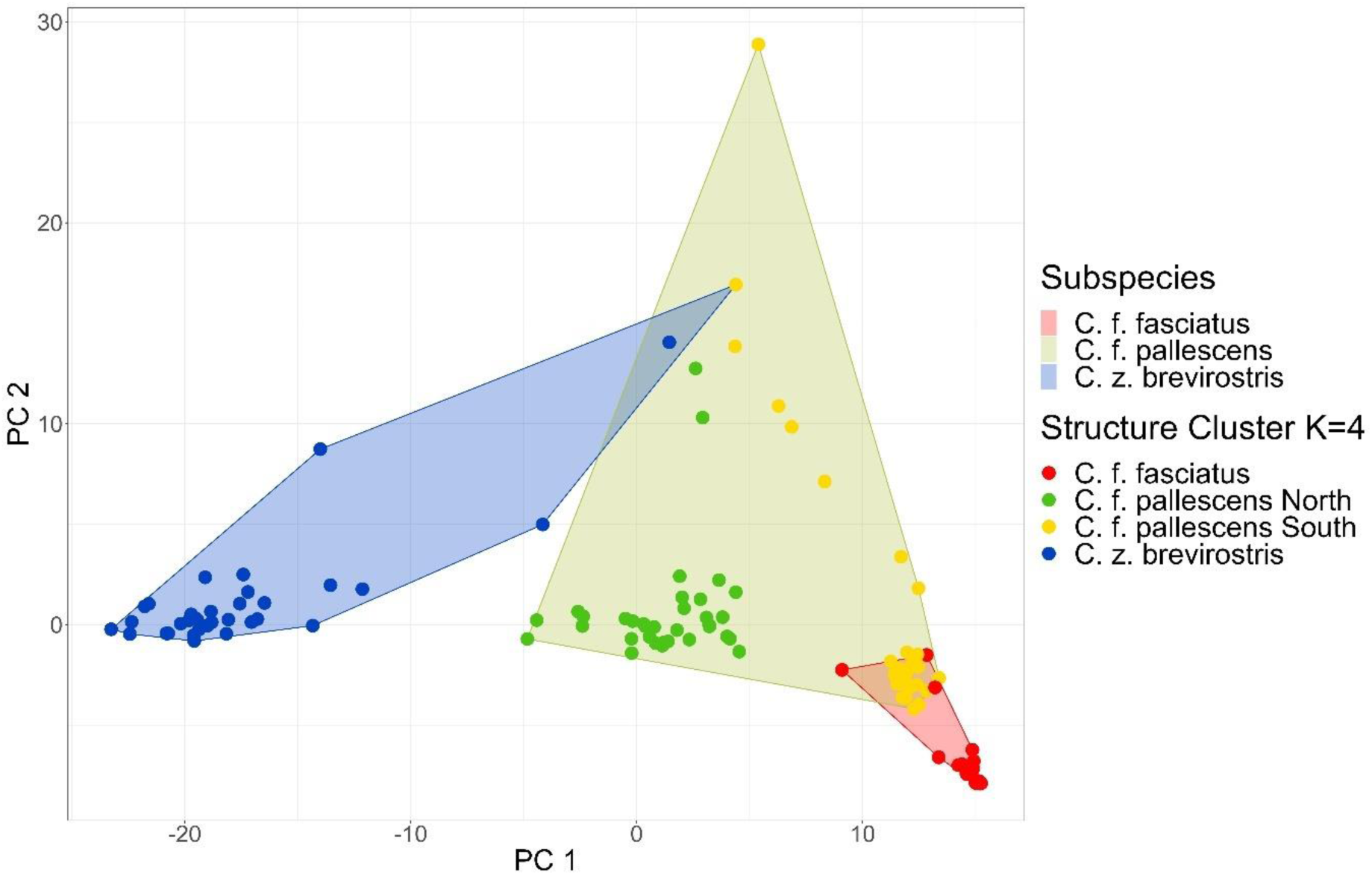
Spatial Principal Component Analyses. The polygons are colored according to the subspecies to which each sample belongs according to Ridgely and Greenfield (2001). The dot colors correspond to the genetic clusters assigned by the analysis with the software Structure when K=4.

STRUCTURE identified K=2 as the most likely number of genetic clusters (Delta K=26,357.04). The second most likely number of genetic clusters was K=3 (Delta K=1,880.47), followed by K=4 (Delta K=351.61)(Fig. 3A). The slope of log-likelihood values of the data against the number of clusters reaches the asymptote at K=2-4 (Fig. S1). We also identified K=2-4 as having the most biological meaning, resembling the clusters observed in the sPCA. Samples from the study region showed a clinal transition from *C. z. brevirostris* from the North to *C. fasciatus* to the south, with admixed individuals falling out in the center (Fig. 3A). The additional DAPC identified four genetic clusters (Fig. 3B) as the optimal model (K=4, BIC=627) and cuts of genetic clusters resembled those from STRUCTURE. Further results from the DAPC showed a sharp decline of BIC for K=2 (641.77) and 3 (630.81).

**Figure 3.**
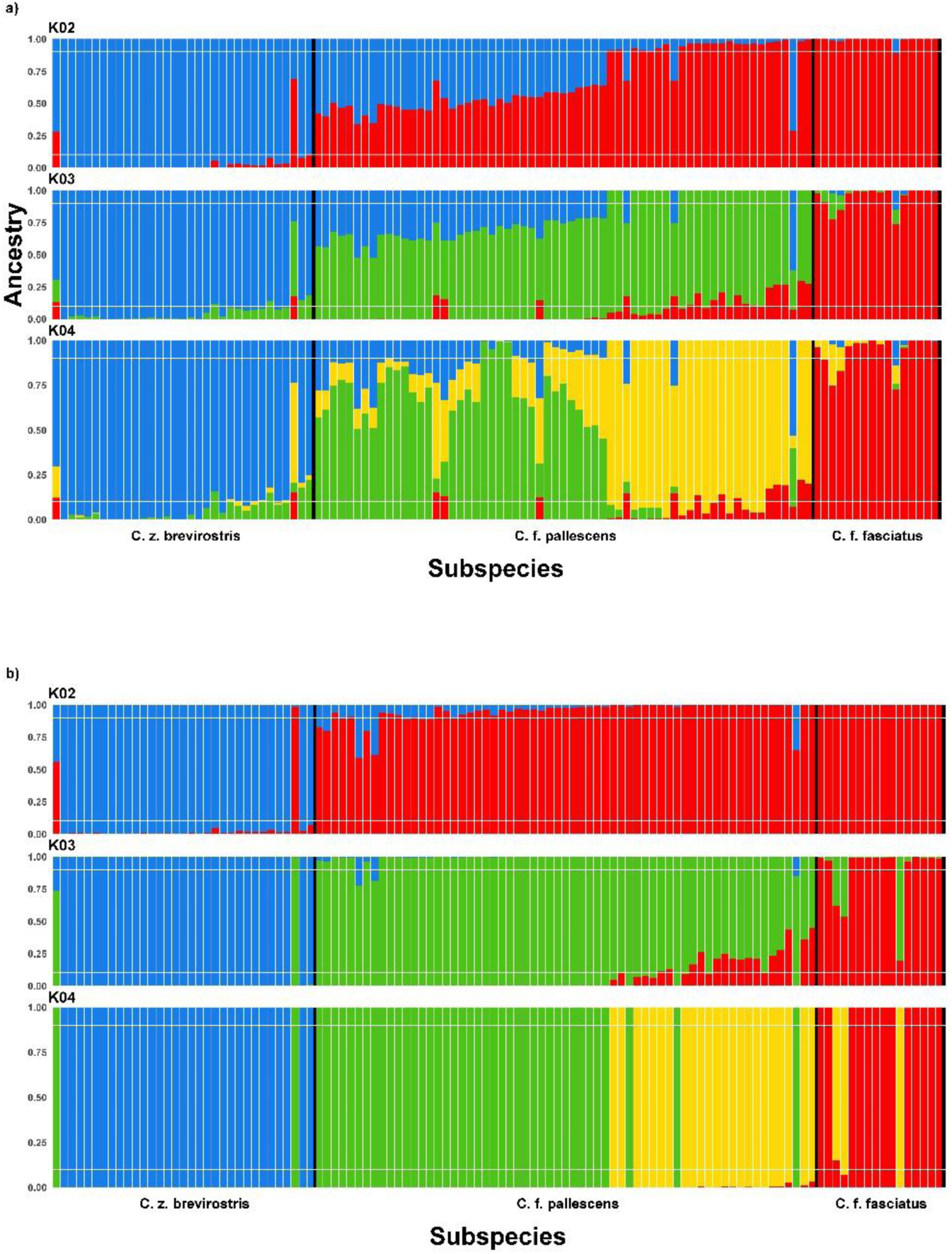
Population Structure. a) Cluster assignment probabilities as estimated by the software STRUCTURE b) DAPC from Adegenet for K=2-4. Samples are ordered by Latitude. Black vertical lines show approximate boundaries among genetic clusters. White horizontal lines show Q=0.9 and 0.1 ancestry probabilities.

At K=2, the distribution of assigned groups resembles the geographical boundary previously reported at the species level with Q values dropping under 0.9 at the sample sites of Chone and as far as Arenillas. When K=3, the assigned group of *C. fasciatus* on the west slope of the Andes matches the geographical distribution of the subspecies *C. f. pallescens*. The occurrence of admixture between *C. z. brevirostris* and *C. f. pallescens* was predominantly observed in the sampling sites spanning from Chone to Manglares Churute. Hereafter, we refer to this genetic cluster as *C. f. pallescens* North. Admixture between *C. f. pallescens* North and *C. f. fasciatus* was identified in the Southwest region of Ecuador and Northwest region of Peru. This genetic cluster is henceforth referred to as *C. f. pallescens* South (Fig. S2).

Hybrid Index HI (proportion of alleles inherited from parental *C. f. fasciatus*) estimated by INTROGRESS showed a cline from the northern-most population of *C. z. brevirostris* (HI=0) to the southern-most population of *C. f. fasciatus* (HI=1) (Fig. 4). HI estimates for individuals across sampling locations showed a gradual increase in the frequency of alleles from parental populations of *C. f. fasciatus*.

**Figure 4.**
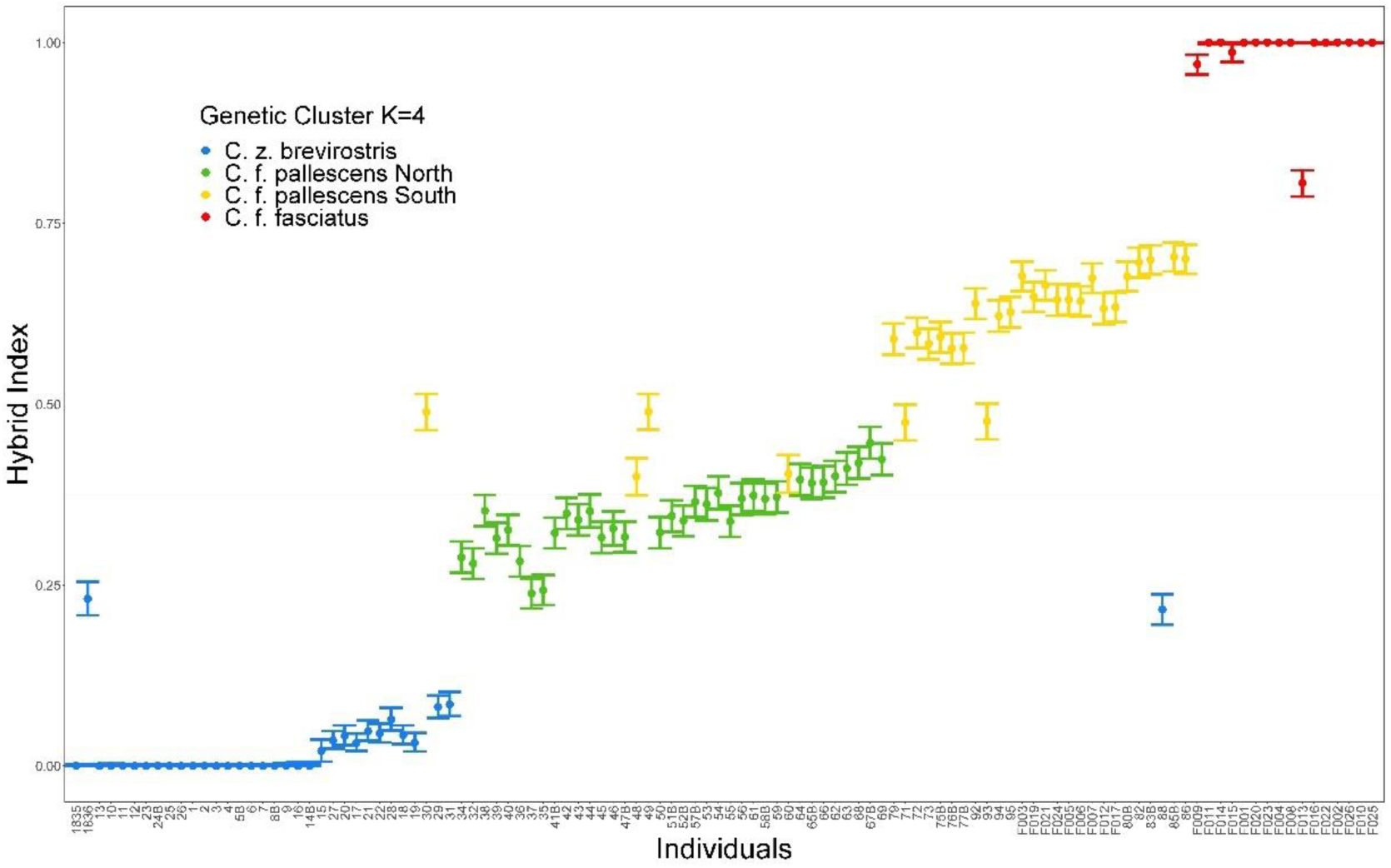
Hybrid Index (HI) per individual as estimated by INTROGRESS R package. Individuals are ordered by Latitude. Colors show different genetic clusters. Samples from parental individuals scored zero at the bottom left and one at the upper right.

### Genetic Diversity

The observed heterozygosity decreased whereas the expected heterozygosity increased when moving towards the southern genetic clusters of *C. f. fasciatus*. *C. f. pallescens* North was the only genetic cluster that showed higher Ho over He (Figure S3 and Table S1).

AMOVA showed that 85.4% (*σ*^2^=0.004, p-value<0.001) of the genetic variation in our data set was within, while 14.6% (*σ*^2^=0.025, p-value<0.001) was among genetic clusters (Table S2). Nei’s Fst showed that the genetic differentiation among distinct genetic clusters was primarily attributed to the differentiation between *C. z. brevirostris* and other genetic clusters. *C. z. brevirostris* showed the highest Nei’s Fst value (0.088) when compared to *C. f. pallescens* South, and the lowest (0.058) when compared to *C. f. fasciatus* (Table S3).

### Isolation by Distance and Environment

Geographical distance was the main factor explaining genetic variation across all analyses, regardless of whether the eastern Andes sampling sites were included, the genetic distance metric used (Nei’s Fst or kinship coefficient), or the statistical method (Mantel test or GDM). The only exception was the Mantel test using Nei’s Fst, which did not reach significance at the α level of 0.001 (r=0.638, p-value=0.0013) (Table 1).

**Table 1.**
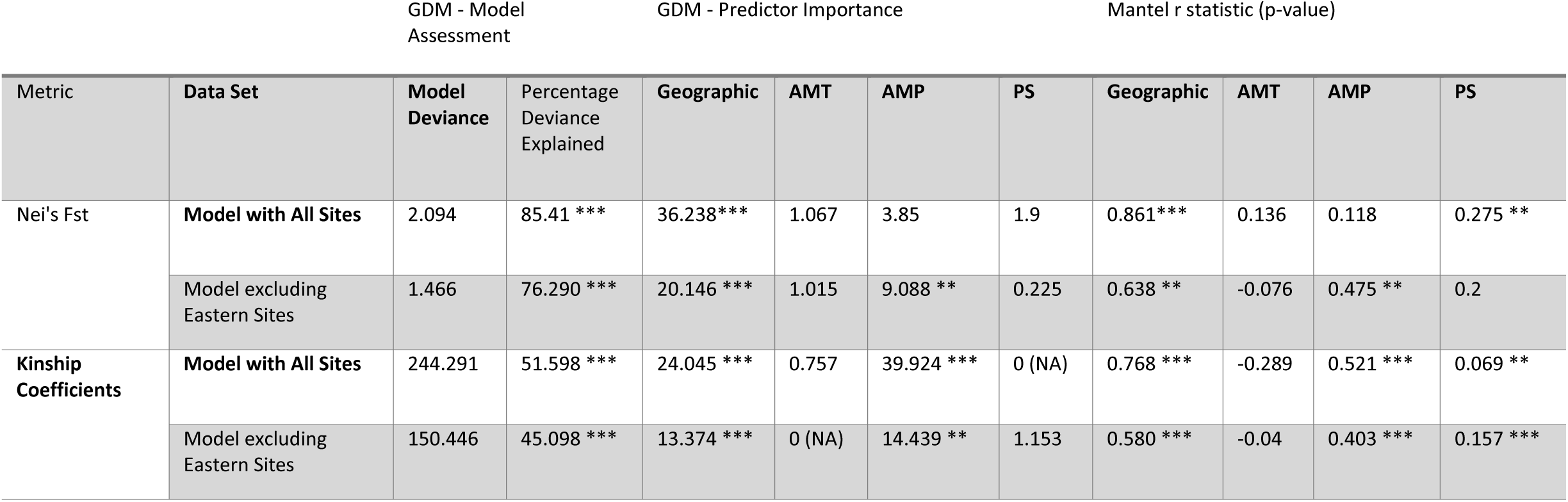
Summary of generalized dissimilarity models (GDMs) and Mantel tests used to explore the effects of geographical and environmental predictors (AMT=annual mean temperature, AMP=annual mean precipitation, and PS=precipitation seasonality) on genetic dissimilarity, as measured by Nei’s Fst distances among sampling sites and normalized kinship coefficient among samples. We performed two sets of analyses, one including and one excluding eastern Andes sampling sites. Predictor importance was measured as the percent decrease in deviance explained between the full model and a model fit with that predictor permuted. Significance is estimated using bootstrapped p-values, with ** indicating a p-value < 0.05 and *** indicating a p-value < 0.001.

| Metric | Data Set | GDM - Model Assessment |  | GDM - Predictor Importance |  |  |  | Mantel r statistic (p-value) |  |  |  |
| --- | --- | --- | --- | --- | --- | --- | --- | --- | --- | --- | --- |
|  |  | Model Deviance | Percentage Deviance Explained | Geographic | AMT | AMP | PS | Geographic | AMT | AMP | PS |
| Nei's Fst | Model with All Sites | 2.094 | 85.41 *** | 36.238*** | 1.067 | 3.85 | 1.9 | 0.861*** | 0.136 | 0.118 | 0.275 ** |
|  | Model excluding Eastern Sites | 1.466 | 76.290 *** | 20.146 *** | 1.015 | 9.088 ** | 0.225 | 0.638 ** | -0.076 | 0.475 ** | 0.2 |
| Kinship Coefficients | Model with All Sites | 244.291 | 51.598 *** | 24.045 *** | 0.757 | 39.924 *** | 0 (NA) | 0.768 *** | -0.289 | 0.521 *** | 0.069 ** |
|  | Model excluding Eastern Sites | 150.446 | 45.098 *** | 13.374 *** | 0 (NA) | 14.439 ** | 1.153 | 0.580 *** | -0.04 | 0.403 *** | 0.157 *** |

AMP was a significant predictor of genetic variation in all models using kinship coefficients, both with and without the eastern Andes sampling sites. It was also significant in the GDM using Nei’s Fst distances, but only when the eastern Andes sampling sites were excluded (predictor importance=9.088, p-value<0.05). PS was only significant in predicting kinship coefficients in the Mantel test without the eastern Andes sampling sites (r=0.157, p-value<0.001). AMT was not significant in any of the models (Table 1).

### Outliers Selection

The MSOD analysis identified seven outlier SNPs (p-value ≤ 0.01) putatively under selection for at least one environmental variable. MSOD identified latitude as the variable with the highest number of associated candidate SNPs (n=6), followed by AMP (n=4) and PS (n=2). Two SNPs were associated only with latitude (snp1219 and snp2476), while one SNP (snp123) was associated with both precipitation variables (AMP and PS). The remaining four SNPs (snp946, snp964, snp2656, and snp3614) were associated with latitude and AMP or PS. MSOD did not detect any SNP associated with AMT. BAYESCENV analysis identified ten putative SNPs as significant for at least one environmental predictor (q-value ≤ 0.01 for γ). Six SNPs were associated with AMT, three with PS and latitude, and two with AMP. One SNP (snp1587) had the highest γ-values for AMT (γ=8.06, q-value<0.01), PS (γ=7.76, q-value<0.01), latitude (γ=7.61, q-value<0.01), and AMP (γ=7.57, q-value<0.01). One SNP (snp4258) was associated with AMT (γ=5.70, q-value<0.01) and latitude (γ=7.26, q-value<0.01). The remaining SNPs were associated with one variable (γ=6.04-2.59, q-value<0.01). No SNPs were identified with both techniques.

### Gene Ontology

A search of the 17 selected outliers yielded 7 genes and 48 GO terms. The GO terms were classified into 15 biological processes, 15 molecular functions, and 18 cellular components. The gene coding for SMG6 nonsense-mediated mRNA decay factor was associated with the most GO terms (Table S4).

### Demographic Scenarios

The best-fit demographic model was isolation with secondary contact and asymmetric gene flow from *C. z. brevirostris* towards *C. f. pallescens* North (For the ancestral Ne = 4.7 × 10^5^, AIC = 23,800.64; Table S5). This model generally had the largest relative likelihood and lowest AIC across the two replicates and ten different ancestral Ne (Fig. S4), except for one of the replicates with ancestral Ne of 2 × 10^5^, 3 × 10^5^, and 7 × 10^5^ where three different models were selected. Ne at the time of divergence in the model of isolation with secondary contact and North-South gene flow was 2.77 × 10^6^ individuals (95% CI 2.77 × 10^6^–2.79 × 10^6^) for *C. z. brevirostris* and 6.87 × 10^4^ (95% CI 5.90 × 10^4^– 6.96 × 10^4^) for *C. f. pallescens* North. Gene flow from the north to the south was 33.38% of Ne (95% CI 30.01% – 44.21%). *C. z. brevirostris* and *C. f. pallescens* diverged around 0.93 million years ago (Ma) (95% CI 8.48 × 10^4^ – 1 × 10^5^ years) (Table S6). Some parameter estimates, specifically migration rates, exhibited a pathological runaway behavior — where the inferred population sizes and epoch durations can degenerate to zero or diverge to infinity — common in SFS-based demographic inference algorithms (Rosen et al. 2018) and therefore should be interpreted with caution.

## DISCUSSION

Here, we analyze the potential contribution of geographical and environmental distances explaining population genetic patterns of a wet-habitat specialist, *C. z. brevirostris*, and a dry-habitat specialist, *C. fasciatus*, along a precipitation gradient in Western Ecuador. We show evidence of genetic structure in western Ecuador and northern Peru. The boundaries of identified groups correspond to the species replacement of *C. z. brevirostris* and *C. f. pallescens* and physical barriers to dispersal. The most likely clusters ranged from K=2-4, corresponding to categories defined by geographic origins, estimated phylogenies, and known physical or ecological constraints. Evidence for IBE due to AMP and IBD was strong across statistical analyses and datasets. We also observed admixture, with a gradual transition between *C. f. pallescens* and *C. z. brevirostris* along the environmental gradient. Genetic differentiation of the two populations of *C. f. pallescens* could be driven by a previously undescribed potential physical barrier near the center of western Ecuador. Lowland habitats in this region may be limited due to the proximity of the Andes to the coastline, limiting dispersal and gene flow, particularly among dry-habitat specialists since much of the habitat is mangrove, wetland and wet forest.

### Population Structure and Admixture Patterns

Although differentiation between western and eastern populations of *C. fasciatus* was expected due to the Andes as a barrier to gene flow, the differentiation of the two western populations of *C. f. pallescens* was not expected. Sampling sites assigned to *C. f. pallescens* in Southwest Ecuador and Northwest Peru formed a discrete group distinct from those in Midwest Ecuador. Heterozygosity patterns coupled with the cline of the ancestry probabilities from STRUCTURE (Fig. 3A), DAPC (Fig. 3B), and HI (Fig. 4) suggested hybridization between *C. f. pallescens* and *C. z. brevirostris* in the contact zone in western Ecuador (Chone).

We believe that admixture events between *C. z. brevisrostris* and *C. f. pallescens* North are suggested to have played a role in the much higher Ho estimated for C. f. pallescens North than found in other cluster (Table S1). The contact zone between hybridizing taxa is expected to exhibit higher levels of heterozygosity (Boca et al. 2020). We hypothesize that reduced gene flow across the Andes may have contributed to the low Ho in *C. f. fasciatus*. Nonetheless, it is important to note that small sample sizes may inflate heterozygosity levels (Schmidt et al. 2021), though utilizing appropriate estimators and a substantial number of bi-allelic markers (>1000), it may be possible to use as few as four individuals (Willing et al. 2012). Thus, it is noteworthy that *C. f. fasciatus* had the lowest sample size (n=12) and Ho values (Table S1). Furthermore, differences in heterozygosity between *C. z. brevirostris* and *C. f. fasciatus* could also be explained by a Wahlund effect, characterized by a decrease in heterozygosity due to a fine-scale population subdivision not accounted for in the sampling (Freeland et al. 2011). Nonetheless, no strong evidence was found to support further population subdivision in our population structure analyses that could lead to the Wahlund effect.

Incomplete lineage sorting (ILS) can also generate genetic diversity patterns comparable to those caused by hybridization (Huerta-Sánchez et al. 2014). ILS refers to the retention and stochastic sorting of ancestral polymorphisms (Maddison et al. 2006). ILS and secondary gene flow can be distinguished when geographic distribution information is available by comparing patterns of genetic diversity between pairs of neighboring and distantly located populations of the different species (Muir and Schlötterer 2005). Gene flow is expected to occur preferentially between neighboring populations, resulting in higher intraspecific genetic diversity and lower interspecific genetic differentiation than between distantly located populations (Petit and Excoffier 2009). In contrast, shared polymorphisms are expected to be distributed evenly across all populations under the ILS scenario (Petit and Excoffier 2009, Zhou et al. 2017). In this study, the progressive increase in the frequencies of alleles as shown by the Hybrid Index (HI, proportion of alleles inherited from parental *C. fasciatus*) differs from the expected allele frequency pattern randomly distributed across two species in ILS. Furthermore, the coalescent-based demographic analysis would identify isolation-with-migration as the best-fit model under ILS (Wang et al. 2019). In contrast, the best model was isolation with secondary contact and asymmetrical gene flow (Table S5).

### Manglares Churute as a Barrier to Gene Flow

The lowlands (from 0 to 800 masl) between the Andes and the coastline constitute a 16km wide corridor near Manglares Churute (a.k.a. Maglares Churute Corridor, MCC). Habitats in the MCC are discontinuous and consist primarily of lentic bodies of water, wetland, second-growth, and evergreen forest with few deciduous and semideciduous remnants (Alava et al. 2007, BirdLife International 2022). In such geographical settings, dispersal between adjacent sites in a one-dimensional stepping-stone model may be limited (Kimura and Weiss 1964). Higher precipitation of the Andean slopes breaks the continuity of arid habitats along the MCC, so dispersal and gene flow may be more difficult for dry-habitat specialists in this region. Along narrow corridors, the effect of IBD restricting gene flow is intensified, particularly for short-range dispersal species (Wright 1943, Kimura and Weiss 1964). Typical dispersal distances for most Troglodytidae remain poorly understood. Current knowledge indicates that Cactus Wrens, for instance, can disperse up to a maximum distance of 26 km and an average of two km (Lynn et al. 2022). Additionally, it is known that cooperative breeding systems may impose constraints on dispersal (Hatchwell 2009). We hypothesize that the *Campylorhynchus* dispersal characteristics, in conjunction with environmental and geographical factors, likely contribute to the restricted gene flow along the MCC.

The barrier to gene flow that ecological settings such as MCC impose on terrestrial lowland species — coupled with anthropogenic threats — might have significant consequences for conservation and evolution (Wagner and Fortin 2013). Other species that show morphometric and plumage differentiation across the MCC could have similar genetic patterns. For example, Necklaced Spinetail (*Synallaxis stictothorax*) has two races: the nominal *stictothorax* occurs north of MCC whereas *maculata* occurs south of the MCC (Ridgely and Greenfield 2001). The same pattern is observed for the nominal race of Collared Antshrike (*Thamnophilus bernardi*), which occurs north of MCC while *piurae* occurs south of the MCC (Ridgely and Greenfield 2001). We suspect that species such as Blackish-headed Spinetail (*Synallaxis tithys*) might exhibit similar genetic differentiation across MCC. If this is correct, it would mean that cryptic biodiversity in the dry forest of west-central Ecuador might need additional conservation attention. We propose that MCC may be an essential barrier to gene flow for lowland dry-habitat specialists and that it should be explored in future studies.

### Isolation by Distance and Environment

We found potential evidence for both IBD and IBE, which was supported by the significant positive relationship between geographic and AMP with genetic distances. Additionally, we detected 17 SNPs associated with climatic factors suggesting potential local adaptations and selection independent of geographic distance. The extent, pattern, and consistency of gene exchange in transitional zones can be explained by both environment-independent (endogenous) and environment-dependent (exogenous) selection (exogenous) (Pyron and Burbrink 2013). Thus, if the selection is exogenous, a clinal genetic pattern such as the one reported in this study (Figs. 3A, 4) may be maintained through differential selection across an environmental gradient, such as a climatic boundary (Haldane 1948, Harrison 2012).

It is important to recognize that although patterns of IBE help identify potential systems for adaptive divergence (Wang and Bradburd 2014), evidence for IBE does not necessarily imply that local adaptations are involved. IBE can arise from several mechanisms other than selection and can be confounded with incipient ecological speciation (Wang and Bradburd 2014). Discerning the relative contribution of geography and environment in shaping genetic diversity remains challenging (Saenz-Agudelo et al. 2015). One major complication in discriminating between these two factors in evolution is that geographical distance and environmental differences are often correlated (Wang and Bradburd 2014, Saenz-Agudelo et al. 2015). Although efforts were made to account for collinearity among predictors in the statistical analyses, it cannot be completely ruled out that collinearity may have affected the reported association in this study.

### Demographic Scenarios and Mechanisms of Introgression

The gradual change of the genomic composition (Figs. 3A, and 4) and the best-fit demographic model (Table S5), suggest introgression from *C. z. brevirostris* into *C. f. pallescens* North. The second best-fit model was isolation with secondary contact and south-north gene flow, implying that gene flow in the opposite direction may also be possible (Table S2). As far as we know, no hybridization involving *C. fasciatus* has been reported previously, but hybridization between *C. albobrunneus* (White-headed Wren) of western Colombia and Panama and *C. z. brevirostris* of Ecuador has been suggested in the north of Ecuador (Ridgely and Greenfield 2001).

One driver of asymmetrical introgression, as we observed here, can be caused by sexual selection through a combination of female choice and/or male-male interactions (Stein and Uy 2006, Martin and Mendelson 2016). In some circumstances, the direction of the introgression is likely to be driven by the sex which determines reproductive choices. For instance, heterospecific female pairing preference for the aggressive golden-collared males in a manakin hybrid zone caused asymmetric introgression of plumage traits into the less aggressive white-collared manakin (Stein and Uy 2006). Similar patterns could also be produced by the lack of female preference for either hetero or conspecifics. For example, introgression skewed toward the Small Tree-Finch (*Camarhynchus parvulus*) in the Galápagos Archipelago was associated with the lack of assortative preference of females of the rarer Medium Tree-Finch (*Camarhynchus pauper*) (Peters et al. 2017). Data on female mating preferences are needed to determine whether female choice drives the introgression of *C. z. brevirostris* into *C. f. pallescens* North.

Interspecific territoriality — a common type of interference competition in animals — is strongly associated with hybridization in birds, implying that reproductive interference favors the maintenance of interspecific territoriality (Cowen et al. 2020, Drury et al. 2020). Interspecific territoriality leads to confrontation between competitors. As a result — regardless of their foraging efficiency — losers in these interactions are frequently excluded from all resources defended by the dominant individual (Gröning and Hochkirch 2008, Grether et al. 2013). Interspecific territoriality could be driving the introgression of *C. z. brevirostris* towards *C. f. pallescens* North if the former is the dominant species. Introgression driven by a dominant species is a pattern found in other species (Pearson and Rohwer 2000, McDonald et al. 2001). Aggressive interference studies are needed to understand the dominant behavior between these species, and whether it is consistent with the directionality of introgression. While the underlying mechanisms behind male-male interactions and female choice are different, they are not mutually exclusive.

When hybrid fitness depends on ecological conditions, fitness consequences of hybridization may vary with environments or fitness components (Parris 2001, Harrison 2012). In such cases, alleles conferring greater fitness under specific ecological conditions may determine the direction of introgression (Coster et al. 2018). Ecological factors might be influencing the introgression of *C. z. brevirostris* into *C. f. pallescens* North if the former has alleles adapted to the climate along the transition zone. An experimental approach to study physiological adaptations and/or genome-wide association studies could help unravel climate as a driving force for introgression between these two species.

The direction of introgression may be influenced by demographic scenarios, such as the highly different population sizes of hybridizing species (Currat et al. 2008, Lepais et al. 2009). According to the “Hubbs principle” — also known as the “desperation theory” — (Hubbs 1955, Randler 2002), birds are more prone to hybridize when the number of individuals in one or both species is limited (Currat et al. 2008, Lepais et al. 2009). The Hubbs principle would predict introgression from *C. f. pallescens* North into *C. z. brevirostris*. In contrast, our results showed that gene flow tends to move towards the larger and more abundant species *C. f. pallescens* North.

## CONCLUSION

Different habitat preferences of *Campylorhynchus z. brevirostris* and *C. f. pallescens*, as well as field observations of potentially admixed individuals, prompted us to search for evidence of isolation-by-distance (IBD) and isolation-by-environment (IBE), and to test hybridization between these species.

We found strong evidence of IBD and some evidence of IBE when we included annual mean precipitation (AMP) in the models. We also found some single-nucleotide polymorphisms (SNPs) that were highly associated with climate. These findings indicate that the genetic variation of *Campylorhynchus* wrens in Western Ecuador and Peru is driven by dispersal limitations as well as potential adaptation linked to variation in precipitation.

Geographical and bioclimatic pressures in Western Ecuador shape genetic variation. As the ranges of these and other species shift under global climate change, it is essential to understand how these pressures shape genetic diversity. By studying the genetic diversity of species across complex bioclimatic landscapes, we can gain insights into local adaptation and the factors that shape genetic variation.

Understanding the impact of climate on genetic diversity is essential for effective conservation strategies in the face of climate change. Genetic diversity is a fundamental objective of conservation biology because it plays a crucial role in facilitating rapid adaptation to environmental changes. By characterizing current ranges and assessing whether species harbor and exchange adaptive genetic variants, we can predict their responses to future climates and inform conservation strategies for wrens and other species with similar distributional patterns.

Populations at niche margins, such as those around the sampling sites of Chone and Manglares Churute, where species ranges approach current and future climate niche limits, likely hold genetic diversity that is critical for adaptation to changing climate. By overlaying genetic monitoring efforts with areas of niche marginality, we can identify where genetic monitoring coincides with anticipated climate change effects on biodiversity.

We do not propose taxonomic changes, but the admixture observed in *C. f. pallescens* suggests that this described subspecies could be a hybrid between *C. z. brevirostris* and *C. fasciatus*, with different degrees of admixture along western Ecuador and northwestern Peru. Further studies including more *Campylorhynchus* species could help to corroborate this hypothesis. Hybridization, despite being a major conservation concern, is also a significant source of novel genetic and phenotypic variation. Hybrid zones provide valuable insights into evolutionary processes. By studying hybrid zones, we can gain a better understanding of the mechanisms that underlie distribution changes, species interaction dynamics, and adaptive introgression. Furthermore, investigating how hybrid zones respond to climate change can provide a more comprehensive understanding of the influence of both abiotic and biotic factors on range limits, as well as how interacting species respond to climate change in the Tumbes-Choco-Magdalena biodiversity hotspot in South America.

Genetic differentiation between populations of *C. f pallescens* across Midwest Ecuador and the disjoint distributions of some dry-habitat specialists suggest the presence of a barrier between Manglares Churute and Arenillas sampling sites. We suggest that this barrier is formed by the proximity of the Andes to the coast and the spatial discontinuity of dry habitats. If this hypothesis is correct, poor dispersers and dry-habitat specialists like the Blackish-headed Spinetail (*Synallaxis tithys*) and the Slaty Becard (*Pachyrhamphus spodiurus*), which are threatened by habitat loss, may experience reduced gene flow across MCC. Reduced gene flow may amplify the synergistic effects of climate change and land use change on these species. These findings could be significant not simply from an evolutionary standpoint but the conservation of species that show similar distribution patterns. Further studies using whole-genome sequencing with greater coverage will allow a more holistic understanding of the influence of both abiotic and biotic factors on range limits and how interacting species respond to climate change.

Understanding the relationship between climate and genetic diversity is crucial for effective conservation strategies in the face of climate change. By characterizing current ranges, assessing adaptive genetic variants, and monitoring genetic diversity, we can predict species’ responses to future climates and inform conservation efforts.

## Acknowledgments

We are grateful for the help of Edith Montalvo, Elise Morton, and the numerous field assistants during the collection of samples. To the Gustavo Orces Museum at the Escuela Politecnica Nacional of Ecuador for including this study in its research management plan. The Ministry of Environment of Ecuador as well as personnel of Reserva Ecológica Arenillas and Refugio de Vida Silvestre El Pambilar for granting the permits for the visits to the sites. We thank the Lovette Lab at The Cornell Lab of Ornithology and particularly Bronwyn G. Butcher for the assistance with the construction of the SNP library and sequencing.

## Authors Contributions

Luis Daniel Montalvo was responsible for the study design, data collection, library construction, sequencing, statistical analyses, interpreting results, writing, and editing of the manuscript. Rebecca T. Kimball contributed to the study design, interpreting results, and editing of the manuscript. James Austin contributed to the interpretation of results, and edition of the document and provide feedback on the statistical analysis. Scott Robinson was largely involved in the study design, interpretation of results, writing, and editing of the manuscript.

## Data Archiving

All sequencing data from this study have been deposited at NCBI Sequence Read Archive under Bioproject accession number PRJNA925654.

---

**Figure S1.**
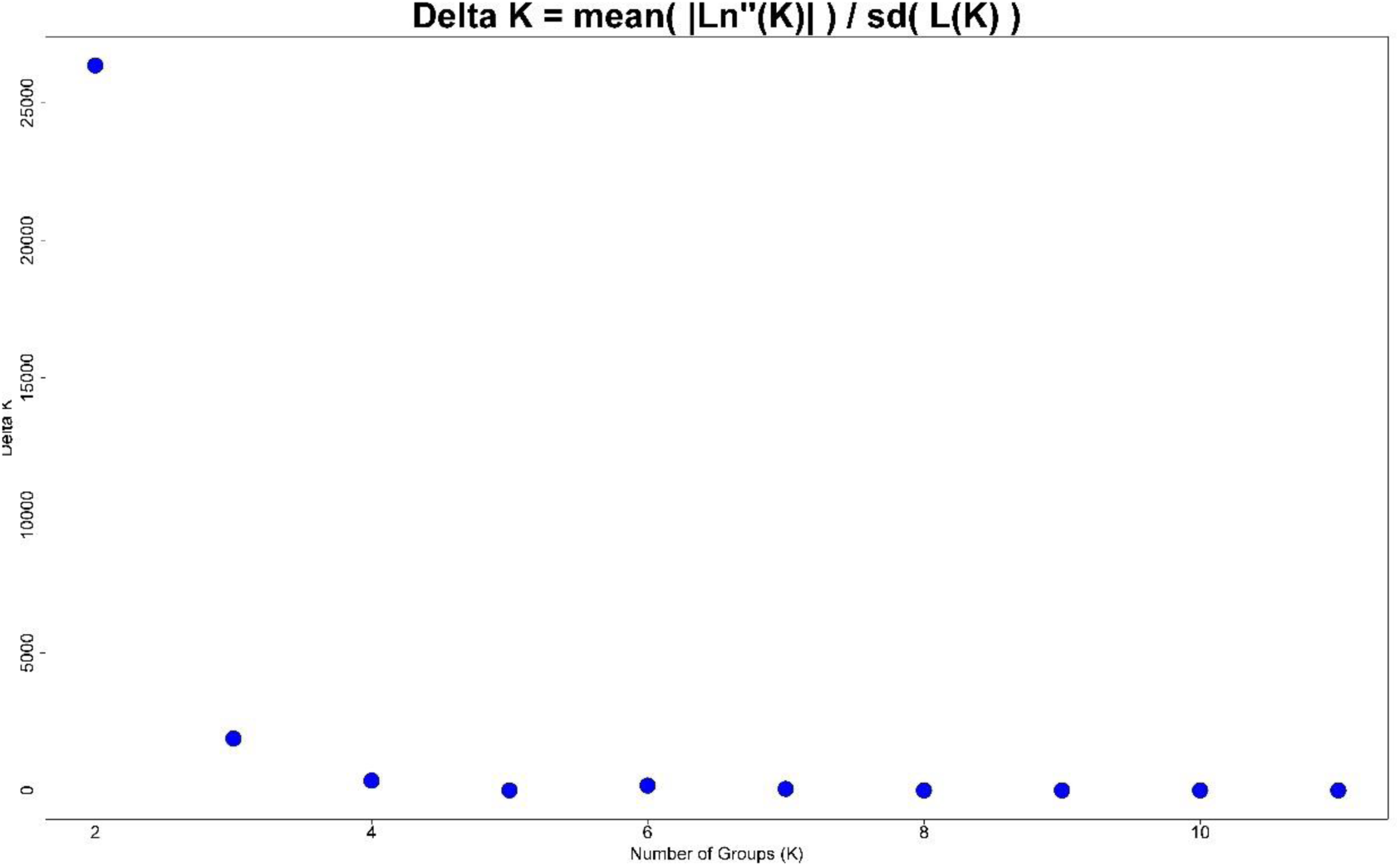
Delta K values for each K as estimated by the software STRUCTURE.

**Figure S2.**
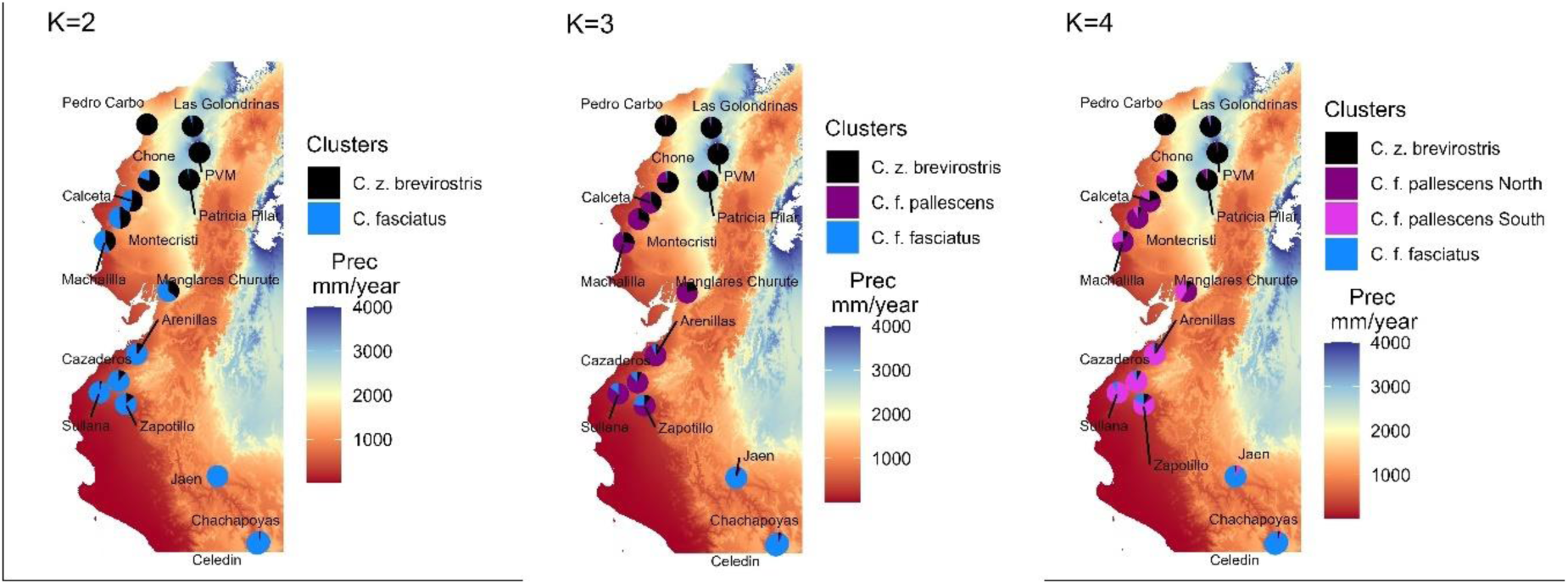
Spatial distribution of mean Q-values per sampling locations along the precipitation gradient in Western Ecuador. Pie charts show the membership probabilities for K = 2 to 4 as estimated for STRUCTURE.

**Figure S3.**
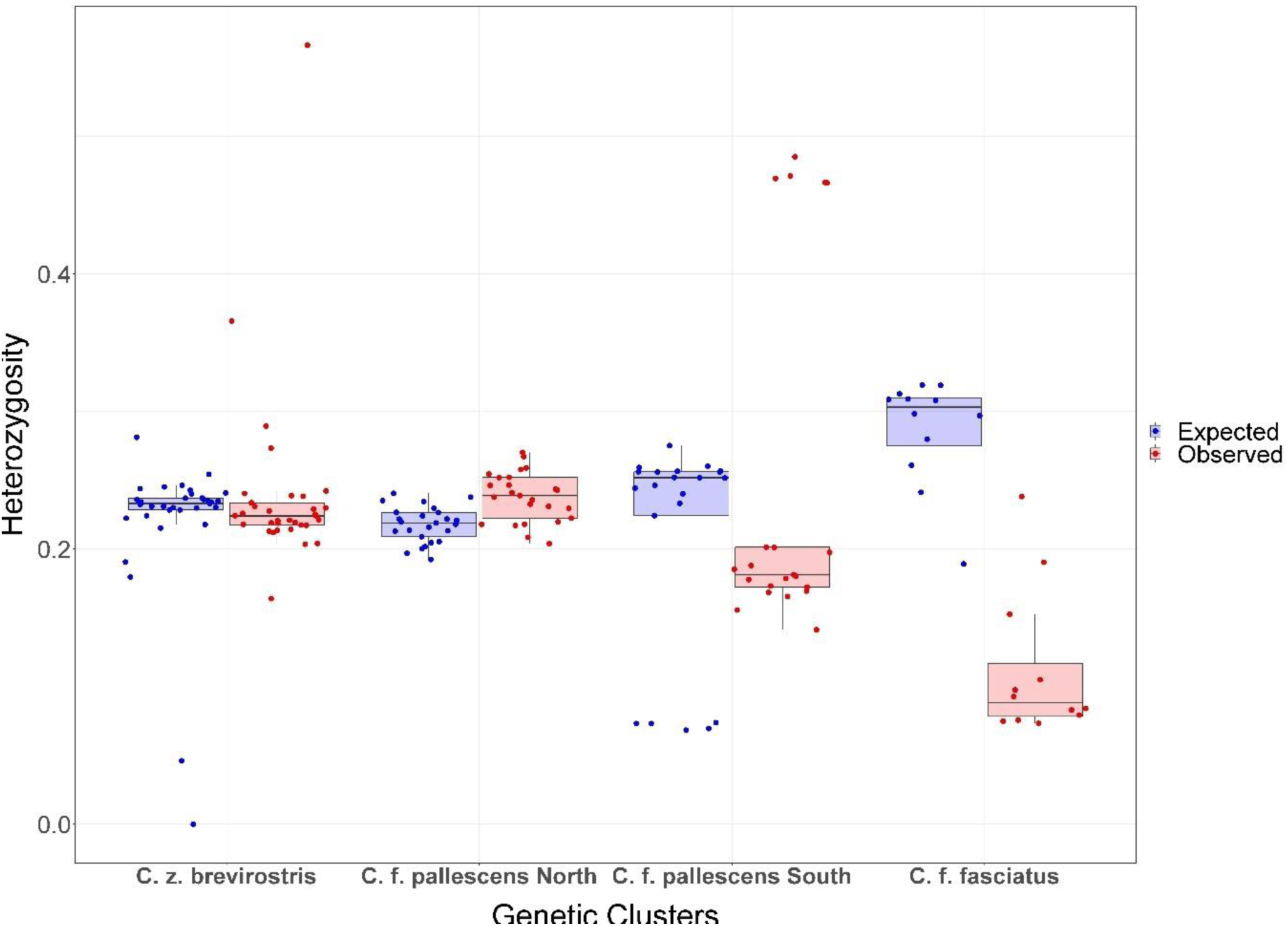
Median of observed and expected heterozygosity and 95 confidence intervals per genetic cluster.

**Figure S4.**
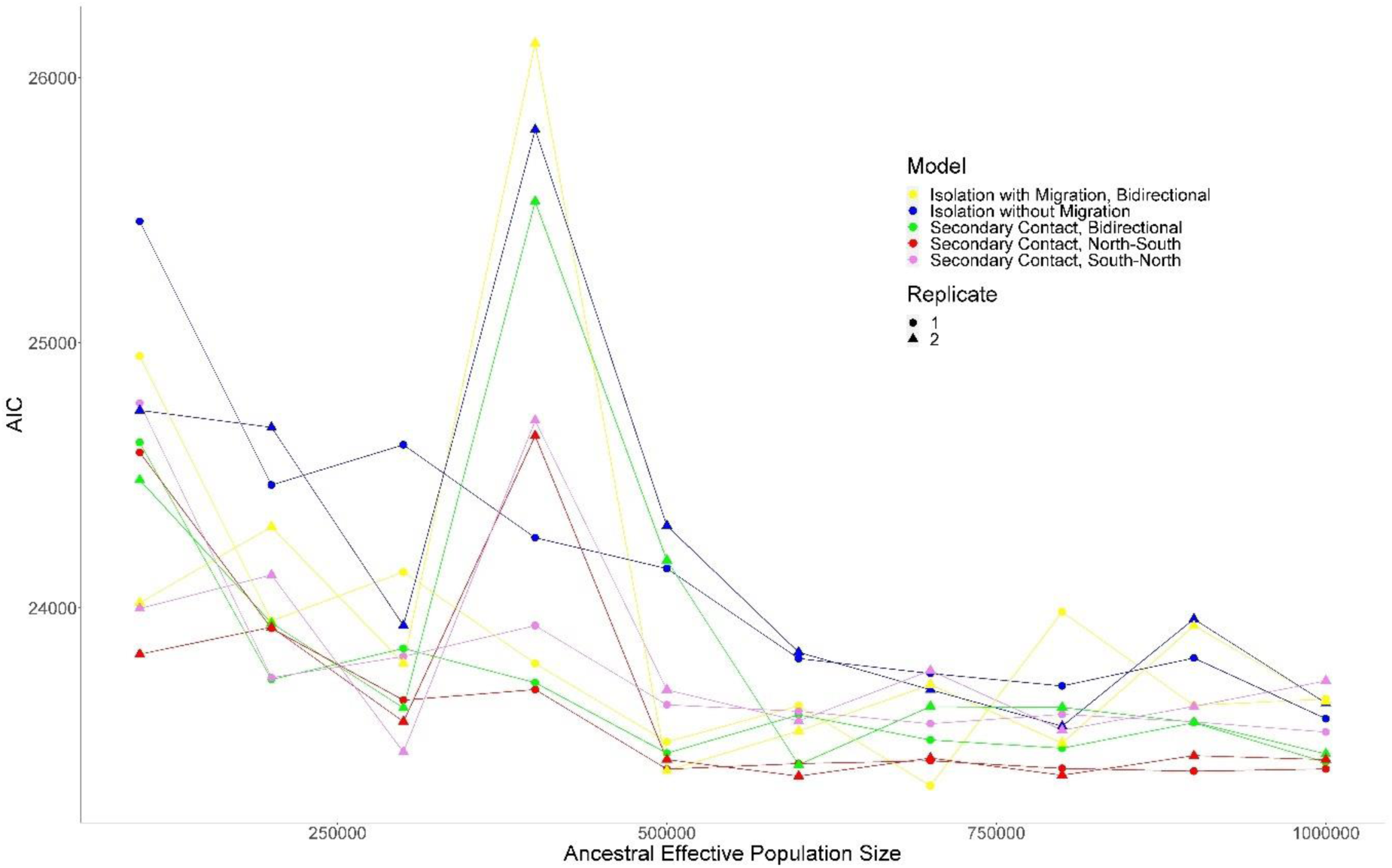
Effects of ten ancestral Ne from 10^4^ to 10^5^ on Akaike information criterion for the five models.

**Table S1.** Genetic clusters assigned by Structure software for K=4, as well as the 2.5%, 25%, 50%, 75%, and 97.5% percentiles of observed heterozygosity (Ho) and expected heterozygosity (He), and inbreeding coefficients (Fis) across individuals for each cluster. The number of samples (N), total number of loci (nLoc), and the number of polymorphic (polyLoc) and monomorphic (monoLoc) loci is reported.

| Genetic Clusters | N | nLoc | polyLoc | monoLoc | Ho Percentiles |  |  |  |  | He Percentiles |  |  |  |  | Fis |
| --- | --- | --- | --- | --- | --- | --- | --- | --- | --- | --- | --- | --- | --- | --- | --- |
|  |  |  |  |  | 2.5th | 25th | 50th | 75th | 97.5th | 2.5th | 25th | 50th | 75th | 97.5th |  |
| C. z. brevirostris | 24 | 4409 | 4254 | 155 | 0.20 | 0.22 | 0.22 | 0.23 | 0.41 | 0.04 | 0.23 | 0.23 | 0.24 | 0.26 | 0.21 |
| C. f. pallescens North | 26 | 4409 | 4322 | 87 | 0.21 | 0.22 | 0.24 | 0.25 | 0.27 | 0.20 | 0.21 | 0.22 | 0.23 | 0.24 | 0.20 |
| C. f. pallescens South | 24 | 4409 | 4341 | 68 | 0.15 | 0.17 | 0.18 | 0.20 | 0.48 | 0.07 | 0.22 | 0.25 | 0.26 | 0.27 | 0.23 |
| C. f. fasciatus | 12 | 4409 | 3918 | 491 | 0.07 | 0.08 | 0.09 | 0.12 | 0.23 | 0.20 | 0.28 | 0.30 | 0.31 | 0.32 | 0.23 |

**Table S2.** Summary of the analysis of molecular variance (AMOVA) within and among genetic clusters as assigned by Structure software for K=4.

|  | SSD | MSD | df | sigma2 | Percentage Variation | P.value |
| --- | --- | --- | --- | --- | --- | --- |
| Among Genetic Clusters | 0.357 | 0.119 | 3.000 | 0.004 | 14.596 | 0.0003 |
| Error | 2.184 | 0.025 | 87.000 | 0.025 | 85.404 |  |
| Total | 2.541 | 0.028 | 90.000 |  |  |  |

**Table S3.** Pairwise Nei’s Fst values among genetic distances as assigned by Structure software for K=4.

| Genetic Clusters | C. z. brevirostris | C. f. pallescens South | C. f. pallescens North | C. f. fasciatus |
| --- | --- | --- | --- | --- |
| C. z. brevirostris |  | 0.088 | 0.078 | 0.058 |
| C. f. pallescens South | 0.088 |  | -0.006 | 0.000 |
| C. f. pallescens North | 0.078 | -0.006 |  | 0.002 |
| C. f. fasciatus | 0.058 | 0.000 | 0.002 |  |

**Table S4.**
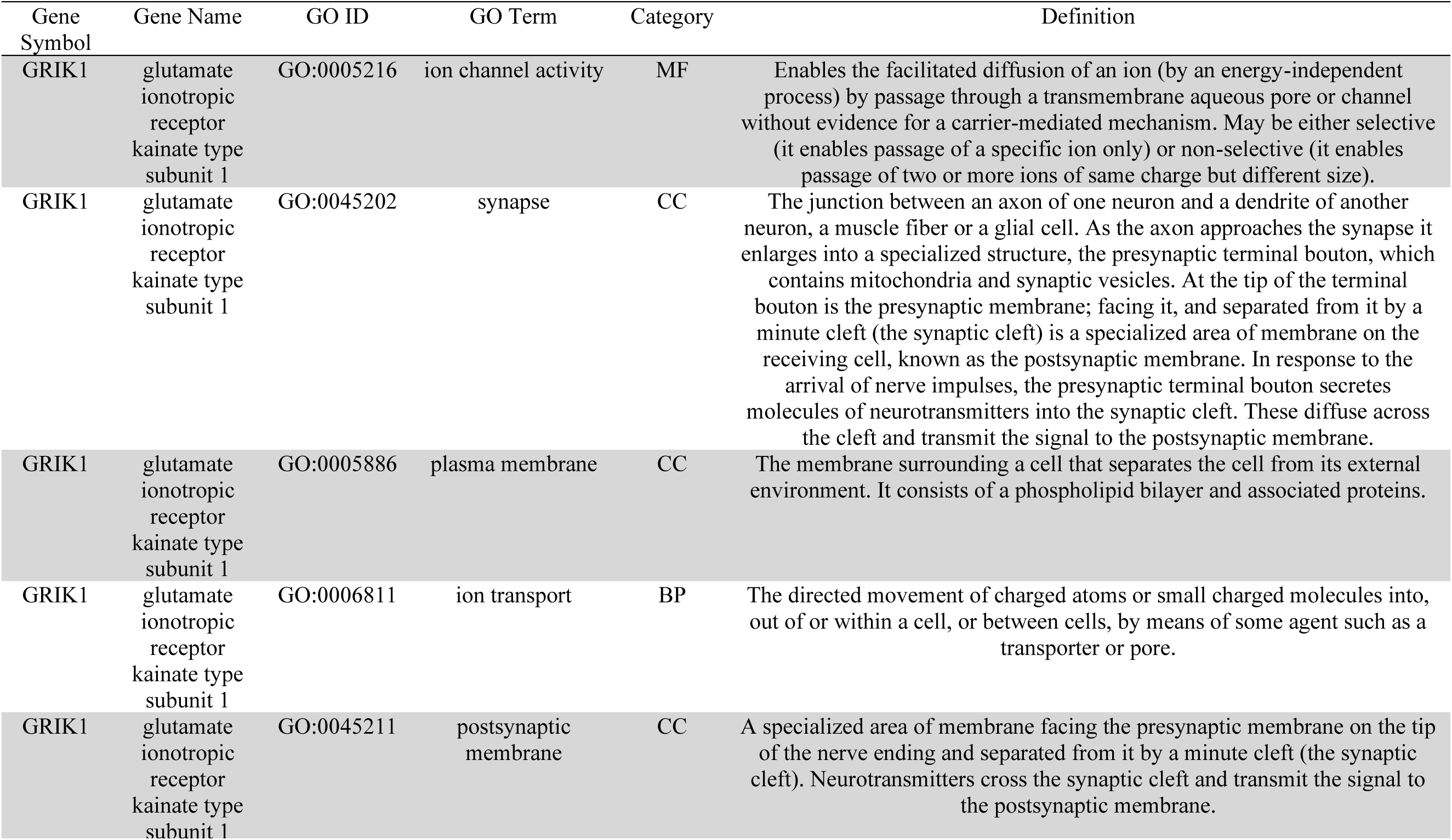
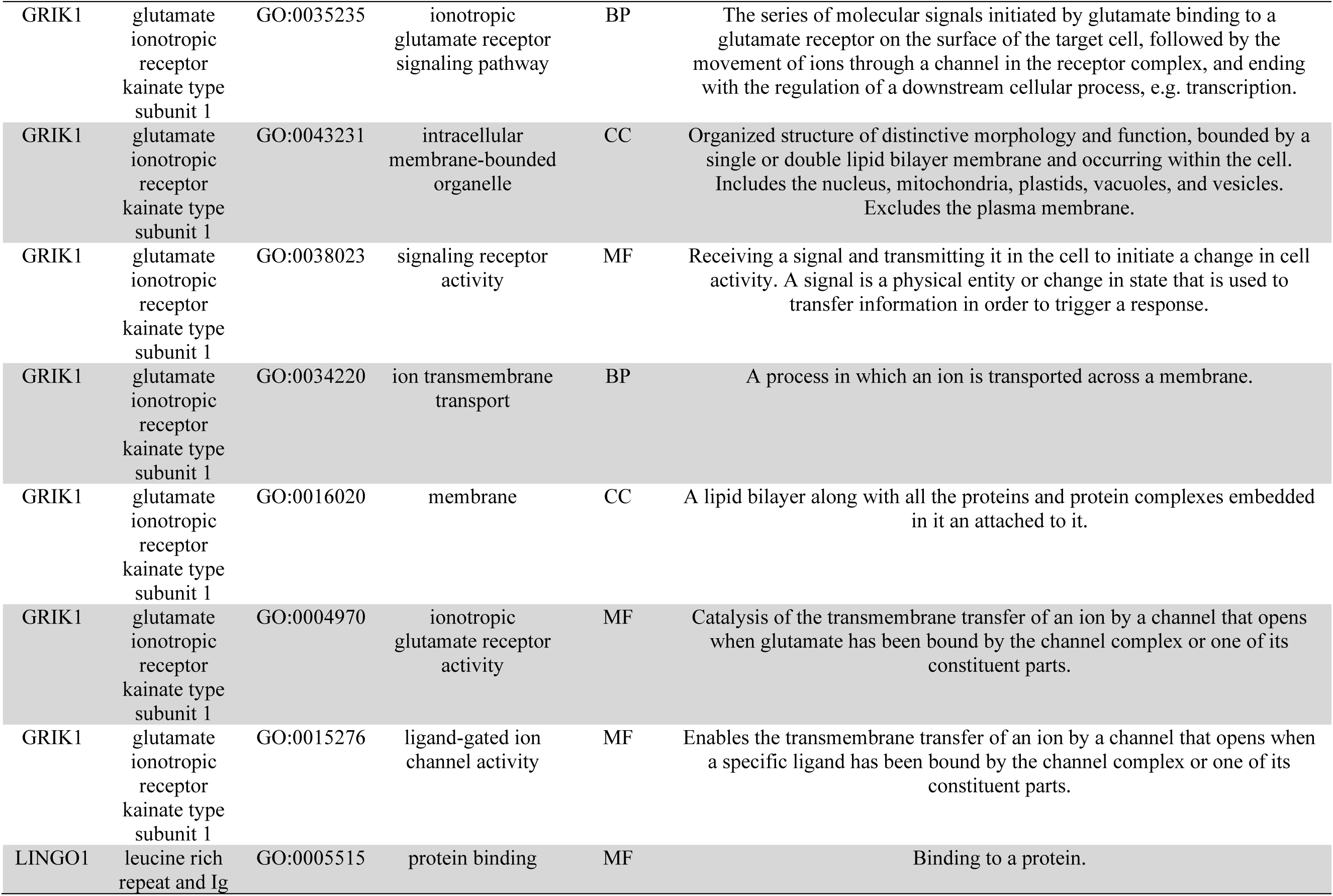
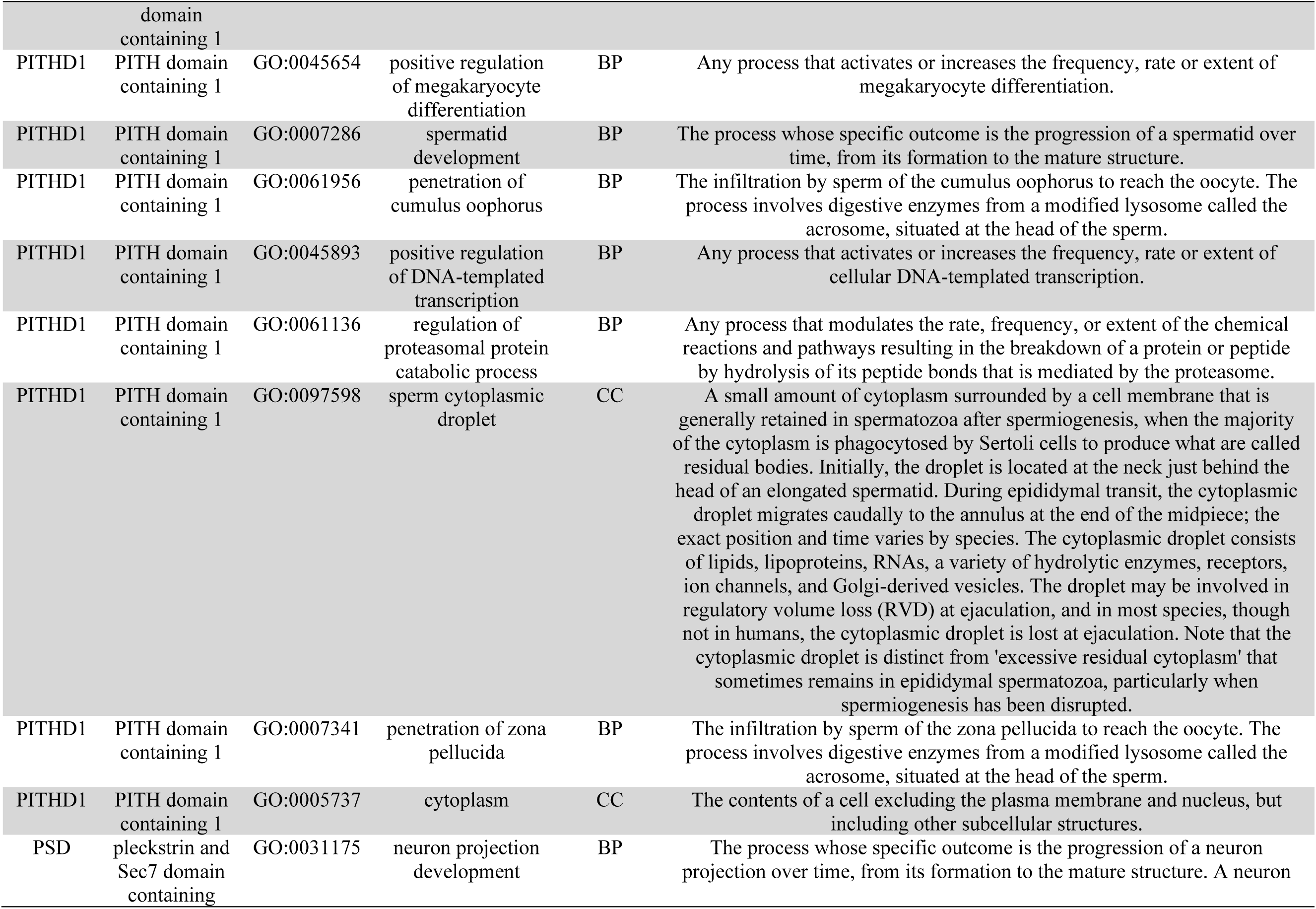
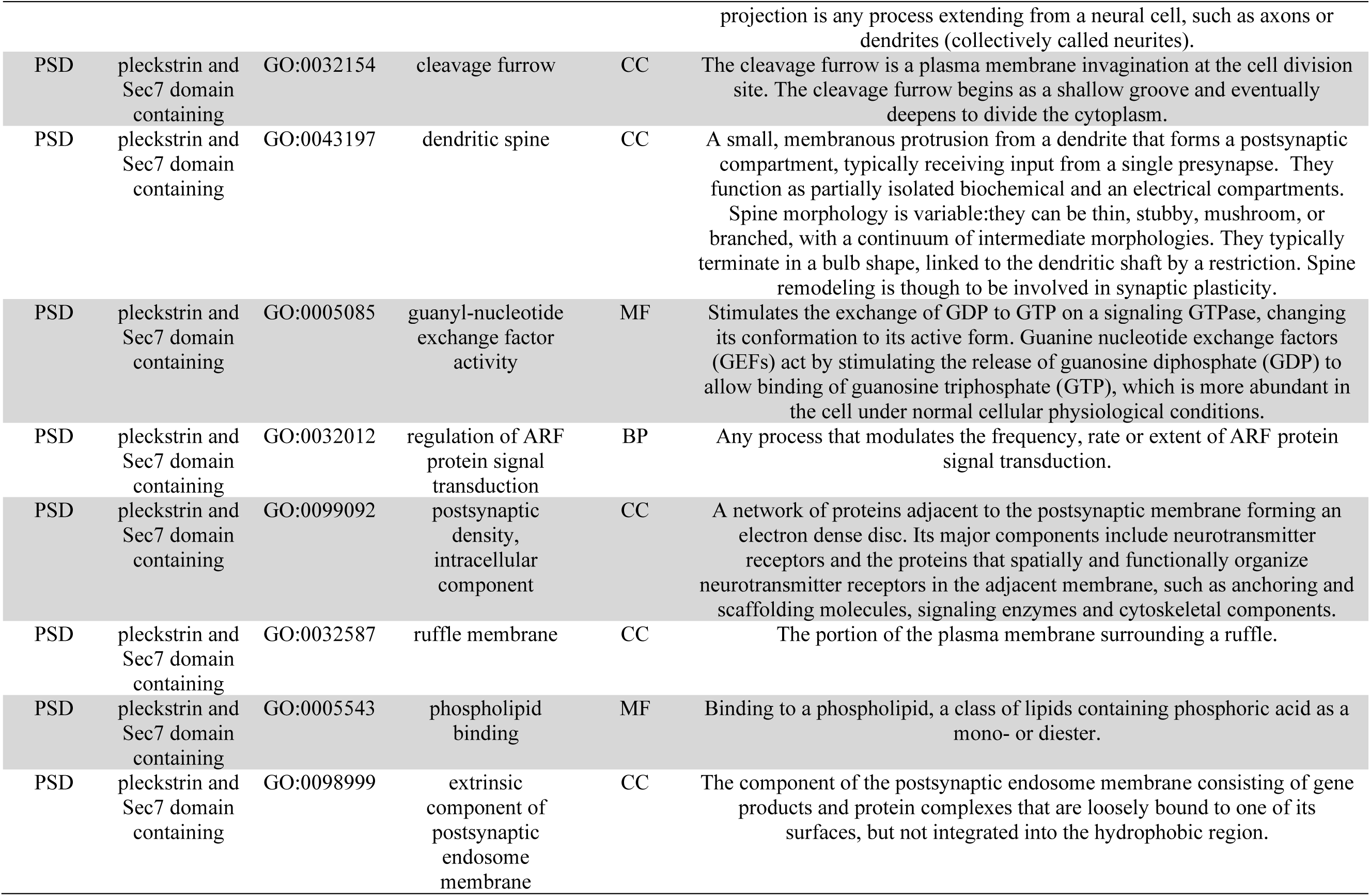
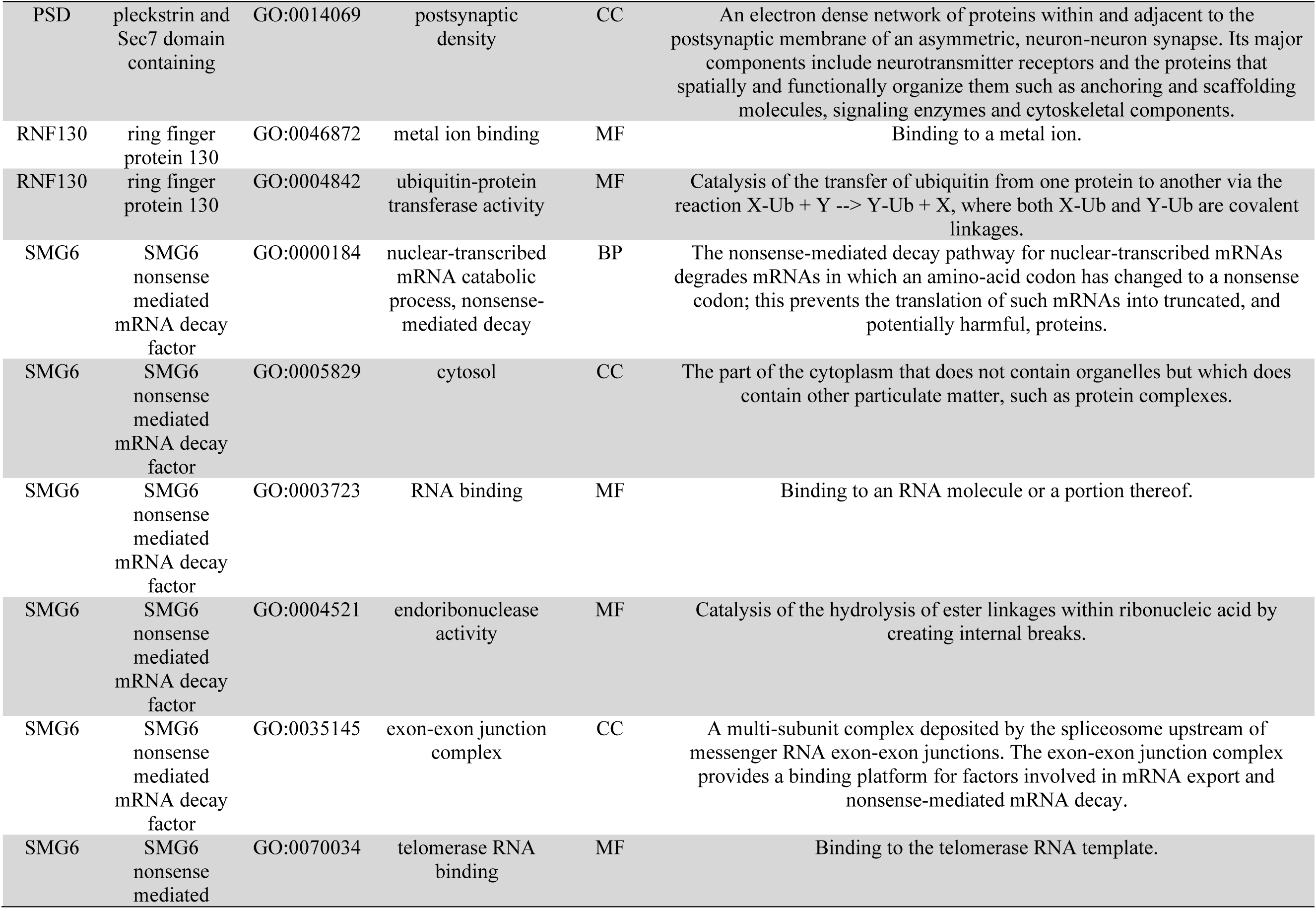
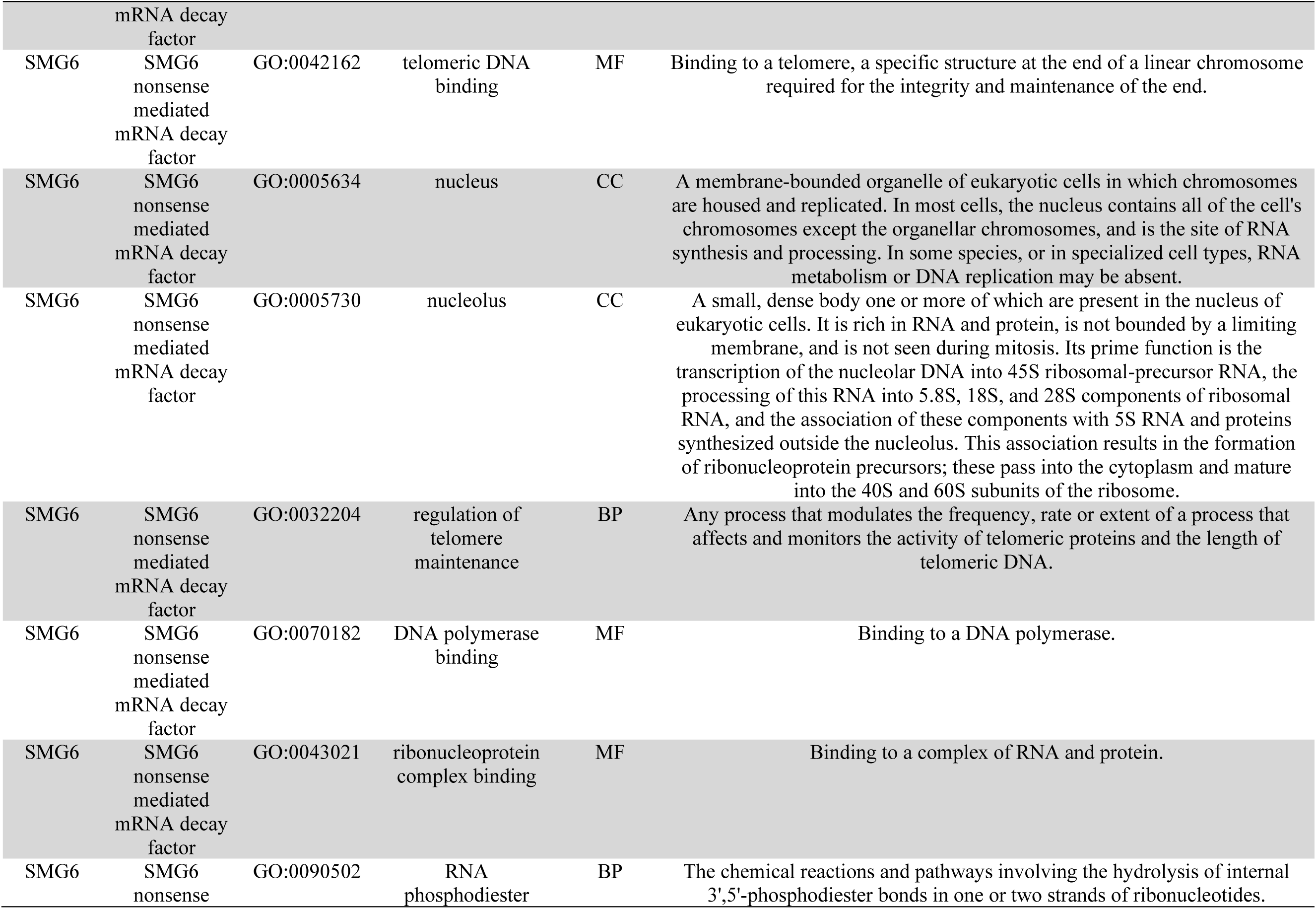
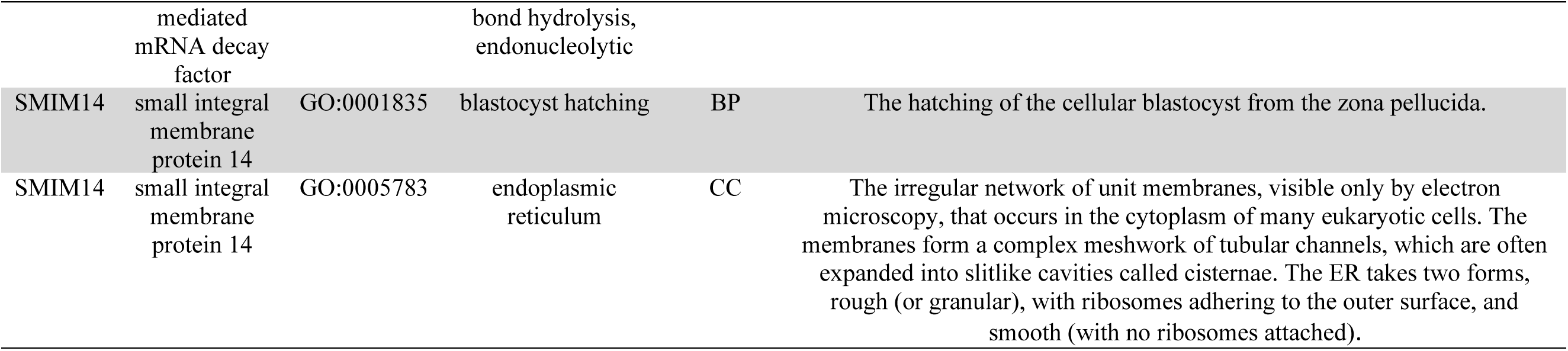
Gene Ontogoly of 17 candidate SNPs associated to climate.BP=biological process, CC=cellular component, and MF=molecular function.

**Table S5.** Model Selection for two-populations models from Momi2. The best model (*) supported asymmetric gene flow between *C. zonatus* towards *C. f. pallescens* North.

| Demographic Scenario | Migration Direction | AIC | delta AIC | AIC weight |
| --- | --- | --- | --- | --- |
| Isolation without Migration | No Migration | 24482.85 | 682.21 | 1.18E-58 |
| Isolation-with-migration | Bidirectional | 24042.54 | 187.92 | 1.56E-41 |
| Secondary Contact | Bidirectional | 23844.24 | 43.60 | 3.41E-10 |
|  | **North-South\*** | **23800.64** | **0** | **1** |
|  | South-North | 24003.52 | 202.87 | 8.85E-45 |

**Table S6.** Descriptive statistics of estimates for effective population size at time of divergence (Necfp_tdiv, Necz_tdiv), time of divergence (tdiv) in years from the present, migration rate (from south to north: msn; from north to south mns) at the time of the migration event or pulse (tmig) from two-populations model between *C. zonatus* (cz) and *C. f. pallescens* North (cfp) analyzed with Momi2. Estimated parameters of best model in *.

| Demographic Scenario | Migration Direction | Parameter | Median | Mean | Lower 95% CI | Upper 95% CI |
| --- | --- | --- | --- | --- | --- | --- |
| Isolation without Migration | **No Migration** | $N_{ecfp\_tdiv}$ | 8.00E+04 | 9.97E+04 | 7.86E+04 | 1.21E+05 |
| | | $N_{ecz\_tdiv}$ | 1.78E+05 | 2.47E+05 | 1.79E+05 | 3.14E+05 |
| | | $tdiv$ | 9.69E+04 | 1.49E+05 | 1.06E+05 | 1.91E+05 |
| Isolation-with-migration | Bidirectional CFP<->CZ | $N_{ecfp\_tdiv}$ | 6.97E+04 | 5.66E+04 | 5.19E+04 | 6.12E+04 |
| | | $N_{ecz\_tdiv}$ | 2.78E+06 | 2.78E+06 | 2.78E+06 | 2.79E+06 |
| | | $m_{sn}$ | 30.40% | 41.95% | 32.34% | 51.55% |
| | | $m_{ns}$ | 38.74% | 42.48% | 34.23% | 50.73% |
| | | $tdiv$ | 9.26E+04 | 9.26E+04 | 8.48E+04 | 1.00E+05 |
| Secondary Contact | Bidirectional CFP<->CZ | $N_{ecfp\_tdiv}$ | 5.78E+04 | 5.70E+04 | 5.17E+04 | 6.22E+04 |
| | | $N_{ecz\_tdiv}$ | 2.77E+06 | 2.78E+06 | 2.77E+06 | 2.79E+06 |
| | | $m_{sn}$ | 27.42% | 35.19% | 26.70% | 43.68% |
| | | $m_{ns}$ | 27.93% | 36.70% | 28.78% | 44.62% |
| | | $tmig_{sn}$ | 1.15E+04 | 1.03E+04 | 8.95E+03 | 1.16E+04 |
| | | $tmig_{ns}$ | 1.21E+04 | 1.08E+04 | 9.70E+03 | 1.20E+04 |
| | | $tdiv$ | 9.26E+04 | 9.26E+04 | 8.48E+04 | 1.00E+05 |
| | North-South CZ->CFP* | $N_{ecfp\_tdiv}$ | **6.87E+04** | **6.43E+04** | **5.90E+04** | **6.96E+04** |
| | | $N_{ecz\_tdiv}$ | **2.77E+06** | **2.78E+06** | **2.77E+06** | **2.79E+06** |
| | | $m_{ns}$ | **33.38%** | **37.11%** | **30.01%** | **44.21%** |
| | | $tmig_{ns}$ | **9.61E+03** | **9.41E+03** | **8.13E+03** | **1.07E+04** |
| | | $tdiv$ | **9.26E+04** | **9.26E+04** | **8.48E+04** | **1.00E+05** |
| | South-North CFP->CZ | $N_{ecfp\_tdiv}$ | 9.06E+04 | 9.59E+04 | 8.31E+04 | 1.09E+05 |
| | | $N_{ecz\_tdiv}$ | 2.77E+06 | 2.77E+06 | 2.77E+06 | 2.78E+06 |
| | | $m_{sn}$ | 25.94% | 33.75% | 25.52% | 41.99% |
| | | $tmig_{sn}$ | 1.15E+04 | 1.05E+04 | 9.26E+03 | 1.18E+04 |
| | | $tdiv$ | 9.26E+04 | 9.26E+04 | 8.48E+04 | 1.00E+05 |

